# Life story of Tunisian durum wheat landraces revealed by their genetic and phenotypic diversity

**DOI:** 10.1101/2020.08.14.251157

**Authors:** Safa Ben Krima, Amine Slim, Sandrine Gélisse, Hajer Kouki, Isabelle Nadaud, Pierre Sourdille, Amor Yahyaoui, Sarrah Ben M’barek, Frédéric Suffert, Thierry C Marcel

## Abstract

Durum wheat (*Triticum turgidum* L. subsp. *durum*) landraces represent a prominent genetic resource for Mediterranean farming systems and breeding programs. Fourteen landraces sampled in Tunisia were genotyped with 9 microsatellite markers and characterized with 15 morphological descriptors, including resistance to the fungal disease *Septoria tritici* blotch (STB). The genetic diversity, nearly was as important within landraces populations (45%) than between populations (54%). It was structured in seven genetic groups and was only partly explained by the variety name or the locality of origin. Populations were also greatly diversified phenotypically (Shannon-Weaver H’=0.54) with traits related to spike and awn colours being the most diversified. Resistance to STB was either qualitative in two populations or with varying degrees of quantitative resistance in the others. A P_st_-F_st_ comparison indicate a local adaptation of the populations. Overall, the genetic structure of Tunisian durum wheat landraces revealed a complex selection trajectory and seed exchanges between farmers.

## INTRODUCTION

The importance of diversity in plant genetic resources used in agriculture and the need for biodiversity conservation is now widely recognized (Maxted & al., 2010). Intra- and interspecific diversity represent a heritage value that is important to preserve because it is the basis for breeders *sensu lato* – from traditional farmers to global breeding companies – to adapt crops to abiotic stresses (*i*.*e*. heterogeneous and changing environments) and provide them with resistance to biotic stresses (*i*.*e*. pests and diseases) (Bellon, 1996). The Green Revolution, which occurred between 1950 and the late 1960s, led to a loss of this diversity. Traditional varieties, thereafter called landraces, were threatened by genetic extinction primarily due to their replacement by modern genetically uniform varieties (Villa & al., 2007). Although humans have historically domesticated and cultivated more than 7,000 species, few high-yielding modern varieties from a limited number of these species constitute nowadays most of the world’s food resources (Perrings, 2018). This evolution to modern production systems promotes the cultivation of some varieties at the expense of traditional and local crops (De luca & al., 2018) and consequently is responsible of a huge intraspecific genetic erosion (Perrings, 2018; Wallace & al, 2019). Moreover, this steady decline of genetic diversity canalizes the evolution of pests and forces to adopt management practices, *e*.*g*. use of pesticides that damage the agro-ecosystem (Conversa & al, 2020). Wild relatives, weedy forms and traditional varieties are especially important for conservation purposes. One of the most threatened components of agricultural plant genetic resources are traditional varieties commonly referred as landraces (Brush, 1997), which constitute the bulk of genetic diversity in domesticated species (Conversa & al., 2020; Poudel & Johnsen, 2009; Villa & al., 2007). Villa & al. (2007) define a landrace as being « a dynamic population of a cultivated plant that has historical origin, distinct identity and lacks formal crop improvement, as well as often being genetically diverse, locally adapted and associated with traditional farming systems ». Landraces are essential heritage from farmer generations at the local and regional scale, as they are associated with traditional farming systems and food trends (Negri, 2003). As a result, they are related to the biological, historical, cultural and socio-economic contexts where they have been grown over generations (Conversa & al, 2020), and are specifically well adapted to the environmental conditions of their cultivation area (*i*.*e*. tolerance to biotic and abiotic stresses) (De Ron & al., 2018; A.C. Zeven, 1998). In conclusion, there is a strong paradox between the loss of landraces being replaced by high yielding modern varieties and the necessity for breeding programs to preserve intraspecific diversity to develop innovative varieties and hybrids (De Ron & al., 2018; Govindaraj & al., 2015; Hammer & Diederichsen, 2009). This statement is particularly relevant to durum wheat.

Durum wheat (*Triticum turgidum* L. ssp. *durum*) is one of the most crucial crops in the Mediterranean countries. It’s a selfing tetraploid species (2n = 4x = 28, AABB) that originated and diversified in the Mediterranean basin (Martínez-Moreno & al., 2020), which is the largest durum wheat producing region worldwide, accounting for about 60% of the total growing area (Royo & al., 2017). This traditional crop is the raw material for the fabrication of local dishes and products including couscous, pasta, several kinds of bread and other semolina products such as bulgur and frike (Belaid 2000; Nazco & al., 2014; Hammami & Sissons, 2020). Durum wheat originated in the Fertile Crescent around 10,000 years B.P. It spread to the western coast of the Mediterranean basin (MacKey, 2005) reaching North Africa around 7,000 years B.P. (Feldman, 2001). During the migration, a combination of natural and farmer’s selection resulted in the development of local durum wheat landraces well adapted to their region of origin and environment. But these durum wheat landraces were recently replaced by improved, genetically uniform and more productive modern varieties also called ‘elite cultivars’ (Soriano & al., 2016). In this context, the National Gene Bank of Tunisia (NGBT) has implemented an ambitious program for the conservation of Tunisian durum wheat landraces both *ex situ* in gene banks and *in situ* on farms. Prospecting activities carried out since 2012 by the NGBT revealed that durum wheat landraces are still cultivated by some farmers in mountainous areas from the North and the Center of Tunisia, under traditional farming systems. These landraces, transmitted by farmers from one generation to the next, are designed by a variety name linked to a historical origin and specific phenotypic characteristics. Previous studies have demonstrated that Tunisian durum wheat landraces are genetically diversified (Medini & al., 2005; Robbana & al., 2019; Slim & al., 2019). Robbana & al (2019) showed a variation of genetic diversity between six durum landraces using Diversity Arrays Technology sequencing (DArTseq). They reported as well a higher level of genetic diversity between landraces than within landraces. Slim & al. (2019) demonstrated for instance a great diversity of 41 Tunisian landraces using 16 molecular markers based on simple sequence repeats (SSRs), also called microsatellites, with clear differentiation between landraces and elite cultivars. They detected also five genetic clusters structuring landraces with a strong North-South stratification. SSRs have been reported to be the most widely used markers to study the genetic diversity in wheat germplasm due to their large distribution in the genome, codominant nature, high polymorphism, good reproducibility and ease of application (Russell & al., 1997; Medini & al., 2005). These multiallelic markers allow to capture higher variability than biallelic markers like Single Nucleotide Polymorphism (SNP), Amplified Fragment Length Polymorphism (AFLP) or DArT markers (Semagn & al., 2014; Targońska & al., 2016; Hurtado & al., 2008).

Several studies have shown that Tunisian durum wheat landraces are agro-morphologically diversified and can be exploited for their large panel of technological properties (Ayed & al., 2010; Nazco & al., 2012; Ayadi & al., 2012; Chamekh & al. 2015; Babay & al. 2019; Bouacha & Rezgui, 2017; Yacoubi & al., 2020). Ayed & al. (2010) showed a huge phenotypic diversity of Tunisian durum wheat landraces for six qualitative traits: seed colour, seed size, glume colour, glume pubescence, spike density and beak length (Ayed & al., 2010). Nazco & al. (2012) highlighted that landraces from the western Mediterranean countries such as Tunisia have heavier grains and higher grain-filling rates than those from the eastern Mediterranean countries. Previous studies established that landraces overstep improved genotypes for agronomic traits such as plant height, biomass, straw yield and also have high grain yield, high nitrogen utilization efficiency, high nitrogen use efficiency, high NADH-dependent glutamate synthase activity and high NADH-dependent glutamate dehydrogenase activity (Ayadi & al., 2012; Chamekh & al. 2015). Additional studies shown that landraces have a high protein content and consequently high physico-chemical technological propriety of semolina and pasta (Babay & al., 2019, Bouacha & Rezgui, 2017). Landraces are known to be sources of increased biomass and thousand kernel weight, both being important traits for adaptation to drought and heat stresses (Yacoubi & al., 2019), but also of genetic resistances to pests, including fungal pathogens. Recently, Huhn & al. (2012) screened a wheat collection including Tunisian durum wheat landraces and revealed that five lines are moderately resistant to Fusarium head blight (Huhn & al., 2012). Ferjaoui & al. (2011, 2015) and Ouaja & al. (2020) identified other lines with resistance to *Septoria tritici* blotch (STB). This foliar disease, caused by the fungal pathogen *Zymoseptoria tritici*, can cause important yield losses in Tunisia (Ferjaoui & al., 2015). Most of the Tunisian durum wheat landraces remain to be genetically and phenotypically characterized, including for their level of resistance to STB disease.

The large majority of studies on durum wheat landraces examine collections of lines belonging to different populations, with only a few or sometimes only one individual per population. It is advisable to extend the population-study to more individuals per population in order to better characterize genetic and phenotypic diversity both between and within populations. To this end, we studied 14 Tunisian durum wheat landraces collected by the NGBT between 2015 and 2017. Some landraces had the same name but were grown in different localities while others were grown by the same farmer but had different names. We thus decided to investigate whether these landraces are different (at the molecular and phenotypic levels) or not and what are their relationships. Concretely, these landraces consisted here and in the whole manuscript as a ‘population’, *i*.*e*. a group of individuals collected at a specific farmer’s field and reported by each farmer to be a landrace. 16 to 51 individuals were randomly selected from each population, which were characterized for: (i) their neutral genetic diversity and structure with SSR markers; (ii) their phenotypic diversity based on agro-morphological characters and their response to STB; and (iii) their phenotypic differentiation (P_st_) comparatively with neutral genetic differentiation (F_st_) in order to determine whether phenotypic differences between populations were due to selection.

## RESULTS

### Identification of hexaploid lineages, historical contaminant of Tunisian durum wheat fields

A first genetic structure analysis of the durum wheat germplasm showed that the lineages from the 14 populations can be divided into 10 genetically distinct groups (strongest ΔK of 14.93) (data not shown). Most genetic groups were composed of lineages coming from different “variety-locality” populations, designed by a “variety” name and its “locality” of origin. At k=10, 33 lineages corresponding to 23 unique multilocus genotypes (MLGs) belong exclusively to three genetic groups genetically close from each other. These lineages stood out for having a characteristic of spikes different from all the others, *i*.*e*. long cylindrical white spikes. These lineages were found in eight out of the 14 populations collected in the North or the Center of Tunisia, *i*.*e*. Roussia Joumine, Mahmoudi Amdoun, Mahmoudi Oued Sbaihia, Mahmoudi El Jouf, Chili El Jouf, Chili Lansarine, Aouija Msaken and Mahmoudi Msaken. They correspond to what is called ‘mule’s tail’ or ‘mare’s tail’ by Tunisian farmers. ‘Mule’s tails’ are undesirable contaminants growing in Tunisian durum wheat fields, whose grains are too “soft” and become flour rather than semolina when milled. The karyotype analysis showed that all mule tails’ lineages suspected to be hexaploid wheat species rather than tetraploid durum wheat carried indeed 42 chromosomes vs. 28 for ‘Karim’ and ‘Mahmoudi-101’ varieties (Figure 1). The highest proportion of ‘mule’s tail’ was found on the two populations Mahmoudi Oued Sbaihia and Mahmoudi El Jouf, both cultivated in the North East of Tunisia (Governorate Zaghouan). Therefore, 33 hexaploid lineages were eliminated from the dataset and further analyses were performed with tetraploid wheat lineages only.

**Figure 1.**
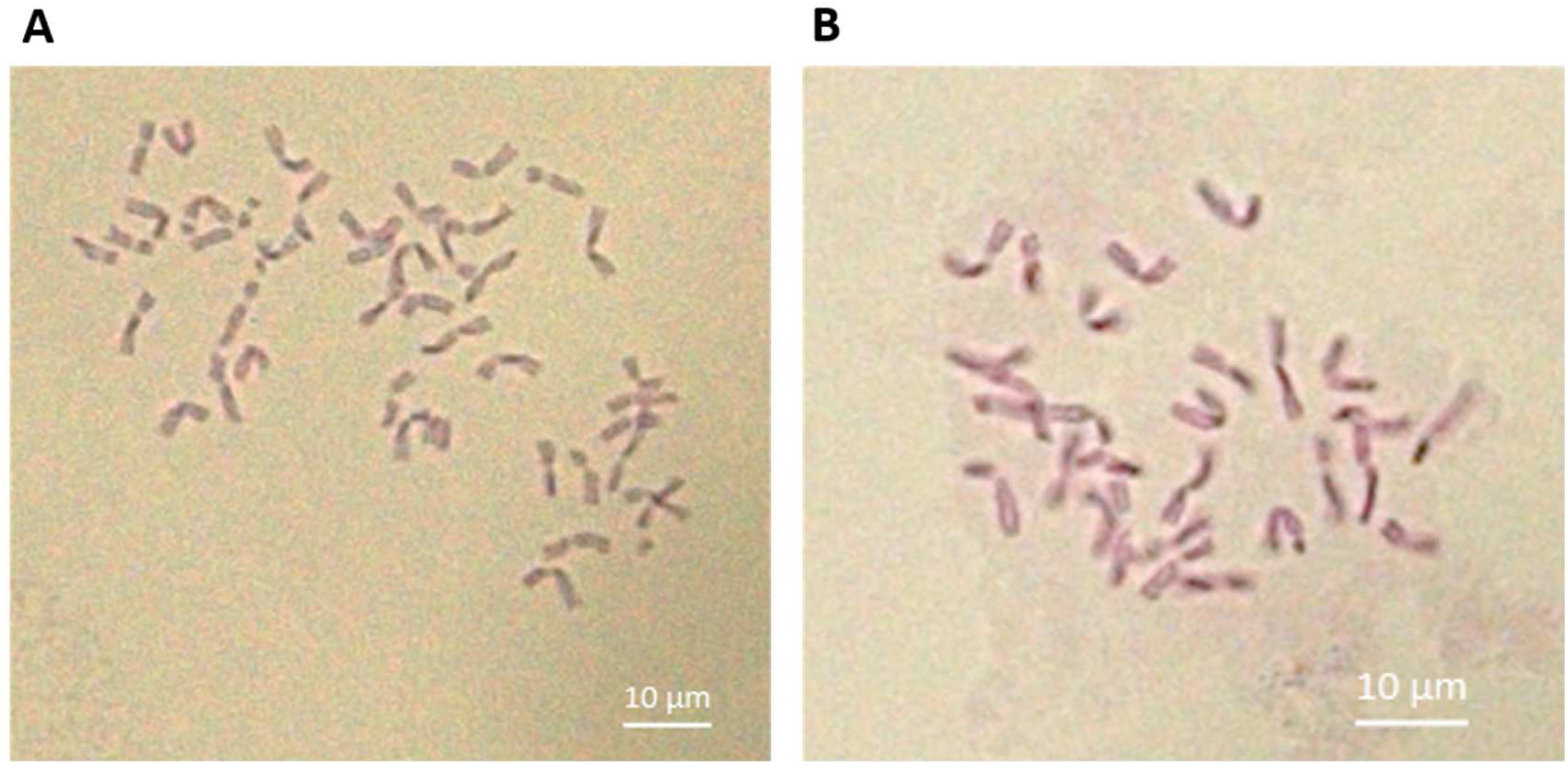
**Karyotypes** of A. a ‘mule’s tail’ lineage carrying 42 chromosomes (hexaploid), and B. the durum wheat cultivar Karim carrying 28 chromosomes (tetraploid).

### Characterization using microsatellite markers

From the collection of 335 lineages of durum wheat, the polymorphic microsatellite (SSR) markers amplified 61 different alleles (Table 1). Size ranges of alleles did not overlap between markers tagged with the same dye. The number of alleles per locus ranged from three for the marker Xgpw2239 to 12 for the marker Xgwm372. In total, 23 private alleles were identified in eight out of the nine loci. The low mean-H_o_ (0.003-0.014) and -H_s_ (0.147-0.317) values revealed a low level of heterozygosity. All markers were highly informative and polymorphic as evidenced from their high PIC value (ranging from 0.503 for Xgpw2239 to 0,966 for Xgwm413), and were characterized by a high Fixation index (0.961-0.986). An analysis of the SSR markers with the BayeScan program revealed no outlier loci. Furthermore, genotype accumulation curves indicated that eight loci only are required to discriminate between individuals in the studied populations (Figure-ESM1). Hence, this multiplex of nine SSR markers is a valuable tool for population genetics studies of durum wheat and genotyping results can be further used for characterizing the diversity of the 14 durum wheat populations.

**Table 1:** Polymorphism of the 9 SSR (microsatellite) markers used to characterize the 14 durum wheat populations: number of alleles (N_a_), number of private alleles (N_ap_), mean observed heterozygosity (H_o_) mean expected heterozygosity (H_s_), fixation index (F_is_) following Nei (1987) expected heterozygosity over all (H_e_), and Polymorphism Information Content (PIC) values.

|  | Xgpw2103 | Xgpw2239 | Xgpw4082 | Xgpw7148 | Xgwm193 | Xgwm285 | Xgwm372 | Xgwm4004 | Xgwm413 |
| --- | --- | --- | --- | --- | --- | --- | --- | --- | --- |
| **N<sub>a</sub>** | 4 | 3 | 6 | 6 | 9 | 8 | 12 | 5 | 8 |
| **N<sub>ap</sub>** | 2 | 0 | 3 | 2 | 5 | 3 | 4 | 1 | 3 |
| **H<sub>o</sub>** | 0.005 | 0.005 | 0.003 | 0.008 | 0.006 | 0.014 | 0.007 | 0.005 | 0.012 |
| **H<sub>s</sub>** | 0.147 | 0.224 | 0.198 | 0.267 | 0.192 | 0.353 | 0.323 | 0.313 | 0.317 |
| **F<sub>is</sub>** | 0.965 | 0.977 | 0.986 | 0.969 | 0.971 | 0.961 | 0.977 | 0.984 | 0.962 |
| **H<sub>e</sub>** | 0.396 | 0.561 | 0.524 | 0.413 | 0.528 | 0.691 | 0.643 | 0.665 | 0.738 |
| **PIC** | 0.322 | 0.423 | 0.466 | 0.387 | 0.492 | 0.639 | 0.618 | 0.600 | 0.704 |

### Distribution of genetic diversity

The AMOVA test determined that there is almost as much inter and intra-population genetic variation (54% of genetic variability was explained by inter-populations variability vs. 45% by diversity within populations; Table 2). As the level of heterozygozity is very low (intra-individual variation around 1%), the studied lineages can be considered fixed after one or two generations of selfing.

**Table 2:** Analysis of Molecular Variance (AMOVA) based on SSR (microsatellite) markers and using the F_st_ measure, for 335 individuals belonging to 14 durum wheat populations.

| Source | df | SS | MS | Estimated variance | Variance (%) |
| --- | --- | --- | --- | --- | --- |
| **Between populations** | 13 | 934.965 | 71.920 | 1.459 | 54% |
| **Within populations** | 321 | 782.721 | 2.438 | 1.204 | 45% |
| **Within individuals** | 335 | 10.500 | 0.031 | 0.031 | 1% |
| **Total** | 669 | 1728.187 |  | 2.694 | 100% |

df: degree of freedom; SS: sums of squares; MS: mean square; variance (%): percentage of total variance contributed by each component.

The genetic diversity indices (genotypic richness; Shannon, Stoddart-Taylor, Simpson and Eveness indexes) for nine SSR markers calculated for each population are given in Table 3. The genotypic richness ranges from 0.107 in the durum wheat population Chili El Jouf to 0.551 in Roussia Joumine. The Shannon index ranges from 0.545 in Chili El Jouf to 2.625 in Roussia Joumine. The Stoddart and Taylor’s index ranges from 1.33 for Chili El Jouf to 11.25 in Roussia Joumine. The Shannon Wiener index is sensitive to genotypic richness in samples of uneven sizes (Grünwald & al., 2003). However, the high positive correlation between the Shannon and Stoddart-Taylor indexes (r=0.86) reinforces the conclusions that can be drawn. The Simpson’s index, measuring the probability that two randomly selected genotypes are different, varies from 0.25 in Chili El Jouf to 0.911 in Roussia Joumine and was also highly correlated with the Shannon index (r=0.96). The Eveness index ranges also from 0.25 in Chili El Jouf to 0.801 in Roussia Joumine, indicating that the MLGs observed in Roussia Joumine are closer to equal abundance than for the other populations. As estimates of genotypic diversity include genotype richness and evenness of distribution of genotypes within the sample (Grünwald & al., 2003), combining results of different evaluated parameters lead us to state that Roussia Joumine stands out as being the most diversified population. Indeed, it has the highest genotypic richness, highest genotypic diversity index and highest Eveness index. On the other hand, Chili El Jouf is the least genetically diverse population.

**Table 3:**
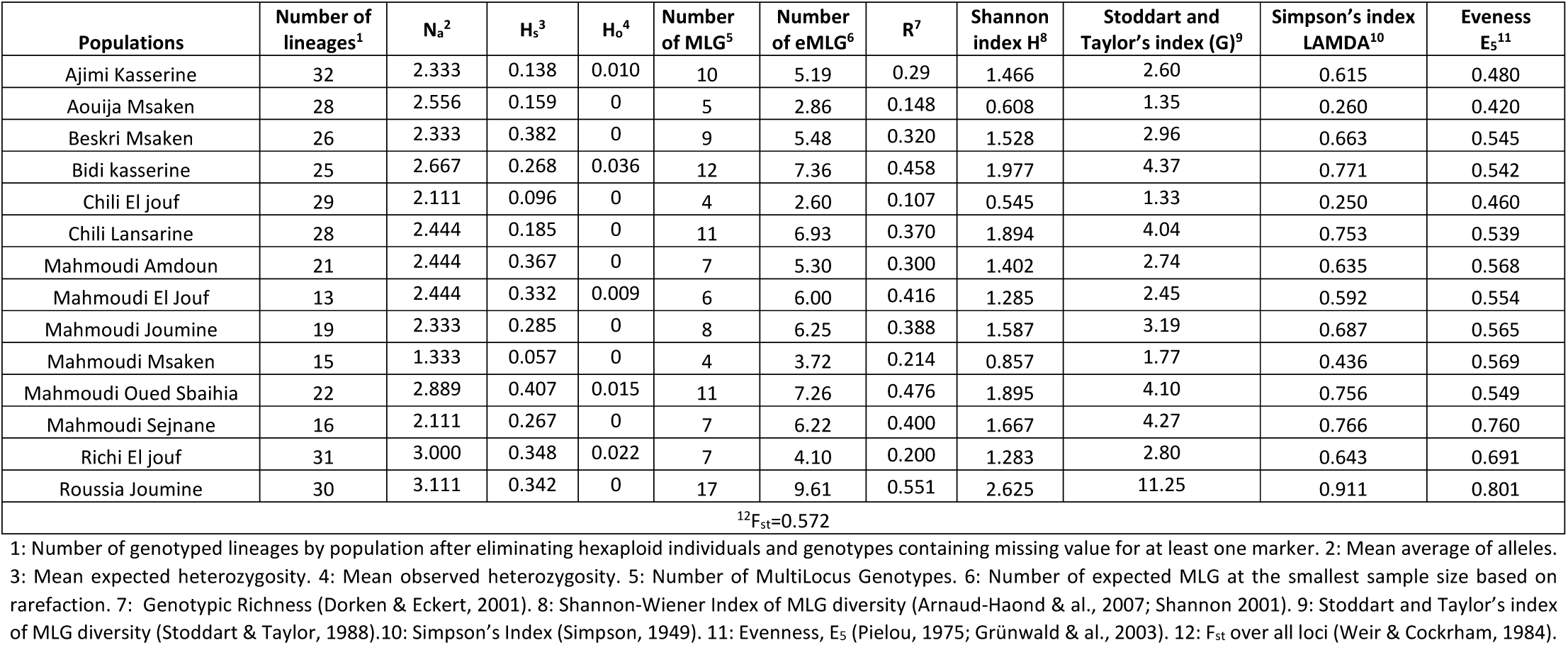
Genetic diversity of the 14 durum wheat populations as evaluated from 9 SSR (microsatellite) markers.

Overall, 118 MLGs were identified from all populations, meaning that 64.8% of lineages are clonal. At equal population sizes, the eMLGs range from 2.6 for Chili El Jouf to 9.6 for Roussia Joumine. Roussia Joumine stands out by its high number of MLGs.

### Genetic structure of Tunisian durum wheat populations

For a number of genetic groups (k) varying from k=3 to k=7, the analysis of the genetic structure of the 14 durum wheat populations highlighted that the strongest ΔK was obtained for k=3 (with ΔK=193.84), k=5 (with ΔK=77.46) and k=7 (with ΔK=44.19) (Figure 2). The number of genetic groups k=7 was chosen because it allows a better description of the genetic structuration of the 14 populations and it maximises the genetic differentiation, *i*.*e*. F_st_, compared to the other partitioning of groups. The seven populations Bidi Kasserine, Ajimi Kasserine, Chili Lansarine, Chili El Jouf, Mahmoudi Msaken, Mahmoudi Sejnane and Roussia Joumine are made of individuals belonging mainly to one genetic group, showing the homogeneity of individuals within these populations. The two populations from Kasserine (collected from the same farmer), Bidi Kasserine and Ajimi Kasserine, are identical, suggesting that they have a similar origin despite their different names. Similarly, individuals from the populations Chili Lansarine and Chili El Jouf also belong to the same genetic group, implying that populations with the same variety name but from different localities aroused from a common origin. The structure of these populations is therefore explained by a combination of effects, including the locality of origin and the variety name.

**Figure 2.**
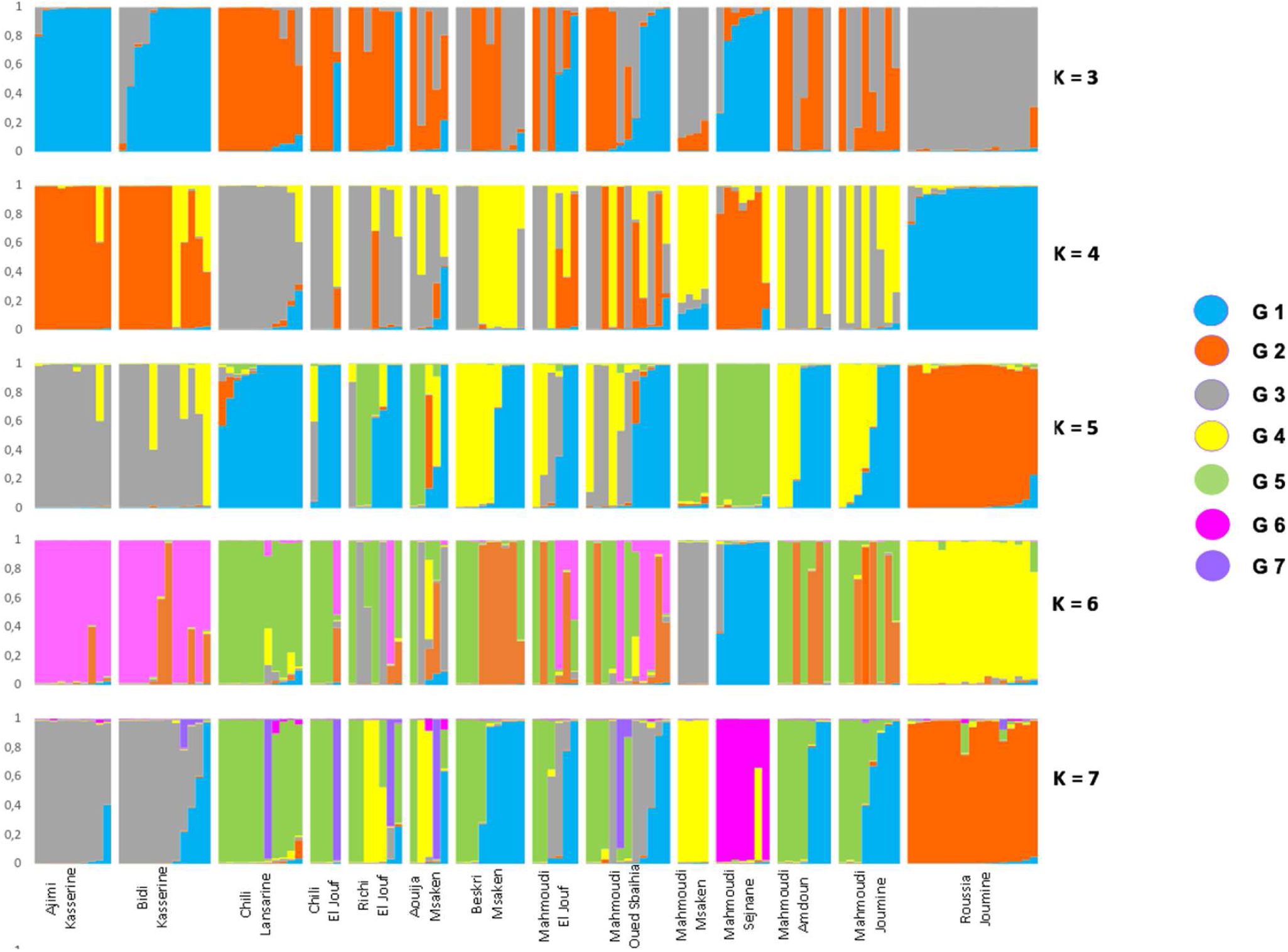
Admixture proportions of the 14 durum wheat populations estimated with STRUCTURE (K=3 to K=7) leading to the identification of different genetic groups (G1 to G7 on the left of the figure). Each vertical bar represents an individual. The colour proportion within each bar represents the posterior probability of assignment of each individual to one of the groups of genetic similarity. The range of assignment probability varied from 0 to 100%.

The minimum spanning network (MSN) visualizes relationships among MLGs and indicates the existence of one MLG of high frequency in nine different populations (Figure 3). The populations sharing this common MLG (MLG.66) are Mahmoudi Oued Sbaihia, Mahmoudi Joumine, Mahmoudi Amdoun, Mahmoudi El Jouf, Richi El Jouf, Aouija Msaken, Beskri Msaken, Chili Lansarine and Chili El Jouf. Other high frequency MLGs were found in two or three populations. Several MLGs detected a single time were highly distant from other MLGs belonging to the same population. MLGs from the population Roussia Joumine were close to each other, and close to MLGs from the two populations Mahmoudi Msaken and Mahmoudi Sejnane. These three populations were distant from the other 11 populations. MLGs belonging to Ajimi Kasserine and Bidi Kasserine were exceptionally close to each other.

**Figure 3.**
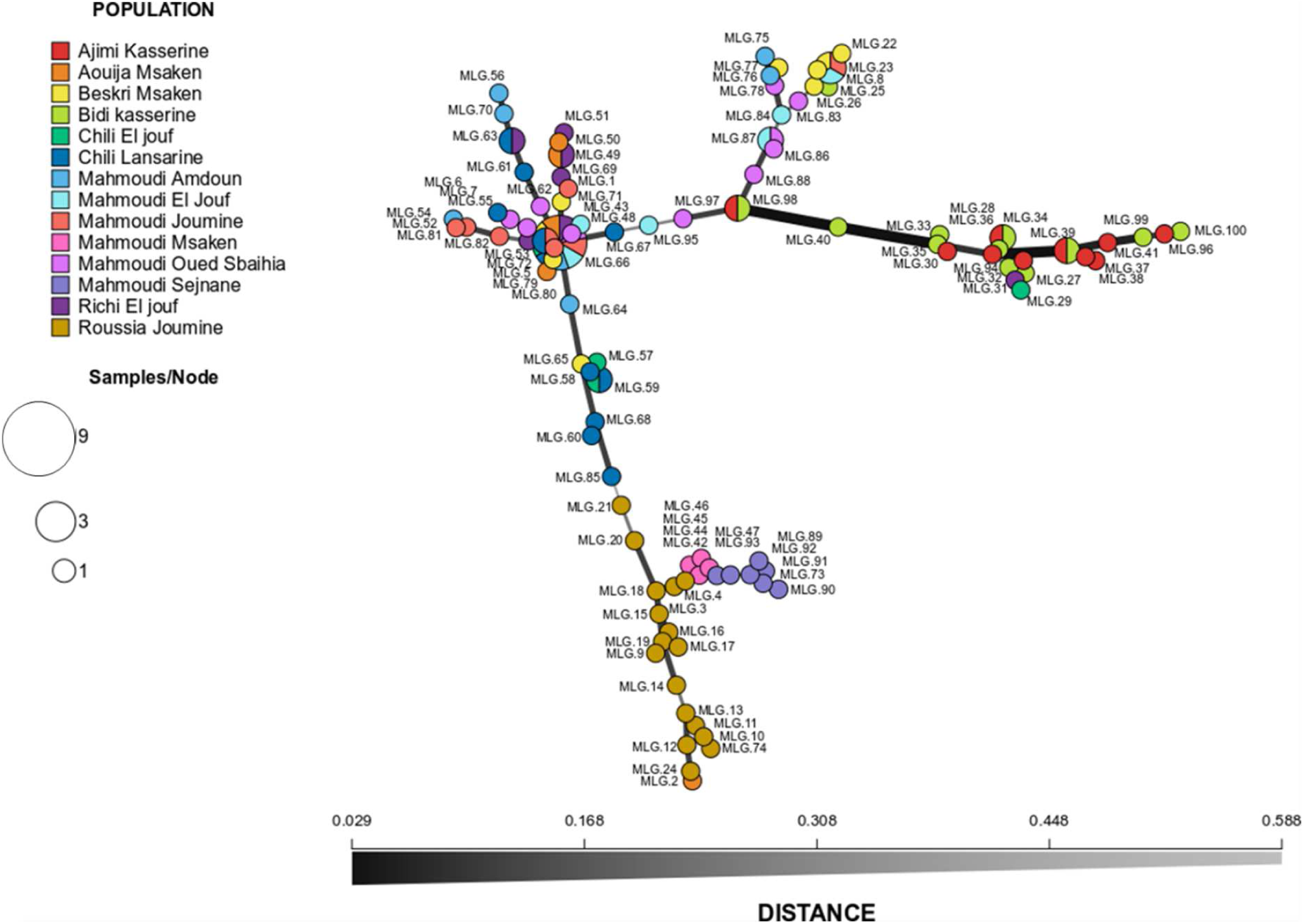
Minimum spanning network between multilocus genotypes (MLGs) of the 14 durum wheat populations. Each node represents a different MLG with the size proportional to the frequency of the MLG. Colours represent the population of origin of each MLG. Edges represent minimum genetic distances between MLGs based on Nei distances (1000 bootstraps) and Neighbor-Joining clustering method. Nodes that are more closely related have thicker and darker edges, whereas nodes that are more distantly related have lighter and thinner edges.

Genetic differentiation between populations was estimated with pairwise F_st_ (F_st_ values between pairs of populations). The matrix of pairwise F_st_ shows values ranging from 0.016 (between Mahmoudi El Jouf and Mahmoudi Oued Sbaihia) to 0.887 (between Ajimi Kasserine and Mahmoudi Msaken), with significant genetic differentiation between the majority of populations (Table 4). In some cases, the genetic differentiation is not significant (5% threshold), for example between the two populations from Kasserine (Bidi and Ajimi varieties), the two Mahmoudi populations from El Jouf and Oued Sbaihia, and the two Mahmoudi populations from Oued Sbaihia and Joumine; while the two Mahmoudi populations from El Jouf and Joumine were significantly different. Similarly, even if significant, the F_st_ value between the two Chili populations from El Jouf and Lansarine is low and very close to the significance threshold. Similarities and differences between populations testify of the history of seed exchanges that took place between farmers. Genetic differentiation between the seven genetic groups determined by STRUCTURE was also calculated using the F_st_ index (Table-ESM1). An individual was considered to belong to a specific genetic group when it has at least 50% affiliation to this group. The matrix of pairwise F_st_ shows values ranging from 0.36 to 0.64 between the seven genetic groups (Table-ESM1) indicating that they are significantly different from each other.

**Table 4:** Matrix of F_st_ between the 14 durum wheat populations.

|  | Ajimi<br>Kasserine | Aouija<br>Msaken | Beskri<br>Msaken | Bidi<br>Kasserine | Chili El<br>Jouf | Chili<br>Lansarine | Mahmoudi<br>Amdoun | Mahmoudi<br>El Jouf | Mahmoudi<br>Joumine | Mahmoudi<br>Msaken | Mahmoudi<br>Oued<br>Sbaihia | Mahmoudi<br>Sejnane | Richi El<br>Jouf |
| --- | --- | --- | --- | --- | --- | --- | --- | --- | --- | --- | --- | --- | --- |
| Aouija Msaken | 0.808* |  |  |  |  |  |  |  |  |  |  |  |  |
| Beskri Msaken | 0.688* | 0.570* |  |  |  |  |  |  |  |  |  |  |  |
| Bidi kasserine | **0.039** | 0.733* | 0.579* |  |  |  |  |  |  |  |  |  |  |
| Chili El Jouf | 0.845* | 0.778* | 0.236* | 0.765* |  |  |  |  |  |  |  |  |  |
| Chili Lansarine | 0.788* | 0.705* | 0.192* | 0.699* | **0.035\*** |  |  |  |  |  |  |  |  |
| Mahmoudi<br>Amdoun | 0.679* | 0.590* | 0.343* | 0.576* | 0.489* | 0.409* |  |  |  |  |  |  |  |
| Mahmoudi El Jouf | 0.722* | 0.618* | **0.047** | 0.591* | 0.149* | **0.088\*** | 0.315* |  |  |  |  |  |  |
| Mahmoudi<br>Joumine | 0.758* | 0.656* | **0.053** | 0.650* | 0.139* | **0.111** | 0.346* | **0.033** |  |  |  |  |  |
| Mahmoudi Msaken | 0.887* | 0.757* | 0.579* | 0.804* | 0.862* | 0.768* | 0.703* | 0.698* | 0.704* |  |  |  |  |
| Mahmoudi Oued<br>Sbaihia | 0.614* | 0.541* | 0.112* | 0.490* | 0.208* | 0.140* | 0.247* | **0.016** | **0.108** | 0.615* |  |  |  |
| Mahmoudi Sejnane | 0.737* | 0.731* | 0.559* | 0.638* | 0.772* | 0.689* | 0.587* | 0.586* | 0.641* | <sup>2</sup>0.800* | 0.527* |  |  |
| Richi El Jouf | 0.680* | 0.270* | 0.233* | 0.591* | 0.354* | 0.287* | 0.315* | 0.188* | 0.253* | 0.575* | 0.178* | 0.570* |  |
| Roussia Joumine | 0.707* | 0.680* | 0.506* | 0.606* | 0.724* | 0.667* | 0.612* | 0.559* | 0.577* | 0.645* | 0.525* | 0.647* | 0.569* |

\*Significance of the $F_{st}$ at a threshold of 5%.

### Phylogenetic relationship between the 14 populations and selection trajectory of Tunisian landraces

Tunisian durum wheat landraces were collected and stored in genebanks since the studies realized by Félicien Boeuf (1875-1961) in the beginning of the 20^th^ century. We studied the relationship between the 14 durum wheat populations collected by the NGBT in 2015 and 2017 and 40 landraces collected between 1911 and 1976 carrying the same variety name (seed samples received from NPGS and NGBT genebanks) (Table-ESM2). The 14 populations, the 40 landraces and individuals from three seed lots of the modern variety ‘Karim’ were genotyped with the nine microsatellite markers described earlier. A neighbour-joining phylogenetic tree was built with microsatellite genotypes (Figure 4). The tree revealed a complex relationship between historical landraces, as samples with the same variety name did not necessarily clustered together. First, the historical landraces called Mahmoudi were mostly divided into two clusters: 14 from the 24 Mahmoudi landraces clustered at the bottom of the tree and were only admixed with the two Bidi landraces (PI534469, 1976) and (PI576736, 1976); 6 of the remaining clustered at the top of the tree and were admixed with one Bidi (Cltr3811, 1912) and one Chili (PI534336, 1976). In this last cluster, the Mahmoudi Msaken population was strongly related to Chili (PI534336, 1976) (bootstrap support of 98.9), and the population Roussia Joumine remains relatively distant from other historical landraces and studied populations. This suggests that Mahmoudi was submitted to at least two different environments, and that in each environment it was confronted to different landraces with which some exchanges could have occurred through open pollination. Second, most of the landraces called Chili were grouped in the same cluster in the middle of the tree, together with few Mahmoudi and Bidi landraces. Among them, landraces Mahmoudi (Cltr15501, 1972) and Bidi (PI157961, 1947) were identical to three Chili landraces. This identity could not have arisen by chance or mutation and this strongly suggests that these two landraces belong to the Chili group and that their names have been badly assigned. This raises the question about the collection and maintenance of accessions in gene banks. Nine of the 14 studied populations also grouped in this cluster with Chili landraces: including Chili Lansarine and Chili El jouf, but also Mahmoudi El Jouf, Mahmoudi Oued Sbaihia, Beskri Msaken, Mahmoudi Joumine, Aouija Msaken, Richi El Jouf and Mahmoudi Amdoun; indicating a strong contribution of Chili historical landraces to the genetic constitution of the studied populations. Third, the populations Ajimi Kasserine and Bidi Kasserine grouped apart from other populations and landraces, and not surprisingly were strongly related to each other (bootstrap support of 100). Finally, in the NGBT and INAT seed lots of ‘Karim’, the same MLG was observed for all individuals with eight of the nine markers, leading us to conclude that these two seed lots are genetically homogeneous; variability of the ninth marker was attributed to a technical bias. For the CRP seed lot, three different MLGs were observed among which the MLG detected in the two other seed lots of ‘Karim’. Thus, this third seed lot corresponds well to ‘Karim’ but is not genetically pure. More surprisingly, Mahmoudi Sejnane was strongly related to Karim (bootstrap support of 98.4), which may indicate that the farmer from Sejnane erroneously considered his seed lot as a landrace.

**Figure 4.**
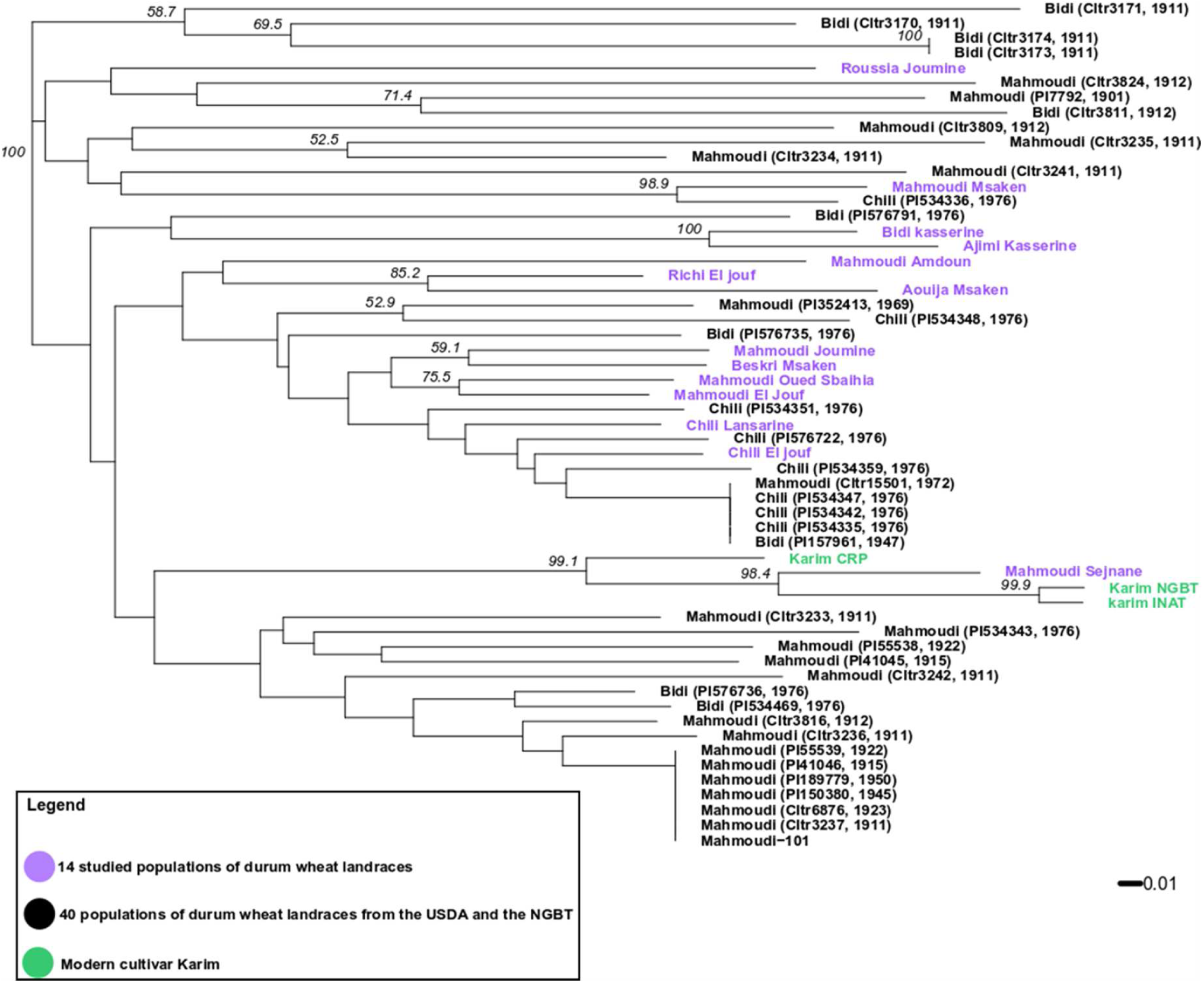
Phylogenetic tree with the 14 studied durum wheat populations (in violet), the 40 landraces collected from the USDA and the NGBT (in black) and three seed lots of the modern cultivar Karim (in green), resulting from Neighbor-Joining cluster analysis based on Edward’s distances (1000 bootstraps). Bootstrap support values expressed in percentages are indicated on the nodes only if there are > 50%.

### Phenotypic diversity of the 14 populations

#### -Shannon Index

The phenotypic diversity was assessed on 273 from the 335 genotyped lineages for the 15 evaluated phenotypic traits (Table ESM3) using the Shannon diversity index (H’) at the population and at the genetic group levels (Table 5). At the population level, the mean H’ for all characterized traits ranged from H’=0.41 in Mahmoudi Msaken to H’=0.66 in Richi El Jouf, making Richi El Jouf the most phenotypically diversified population. The spike colour (SC) was the most polymorphic trait over the 14 populations (mean H’=1.06), with spikes being mostly slightly coloured. The seed shape (SS) was the least diversified trait over all populations (mean H’=0.08), with the majority of populations having semi-elongated seeds.

**Table 5.**
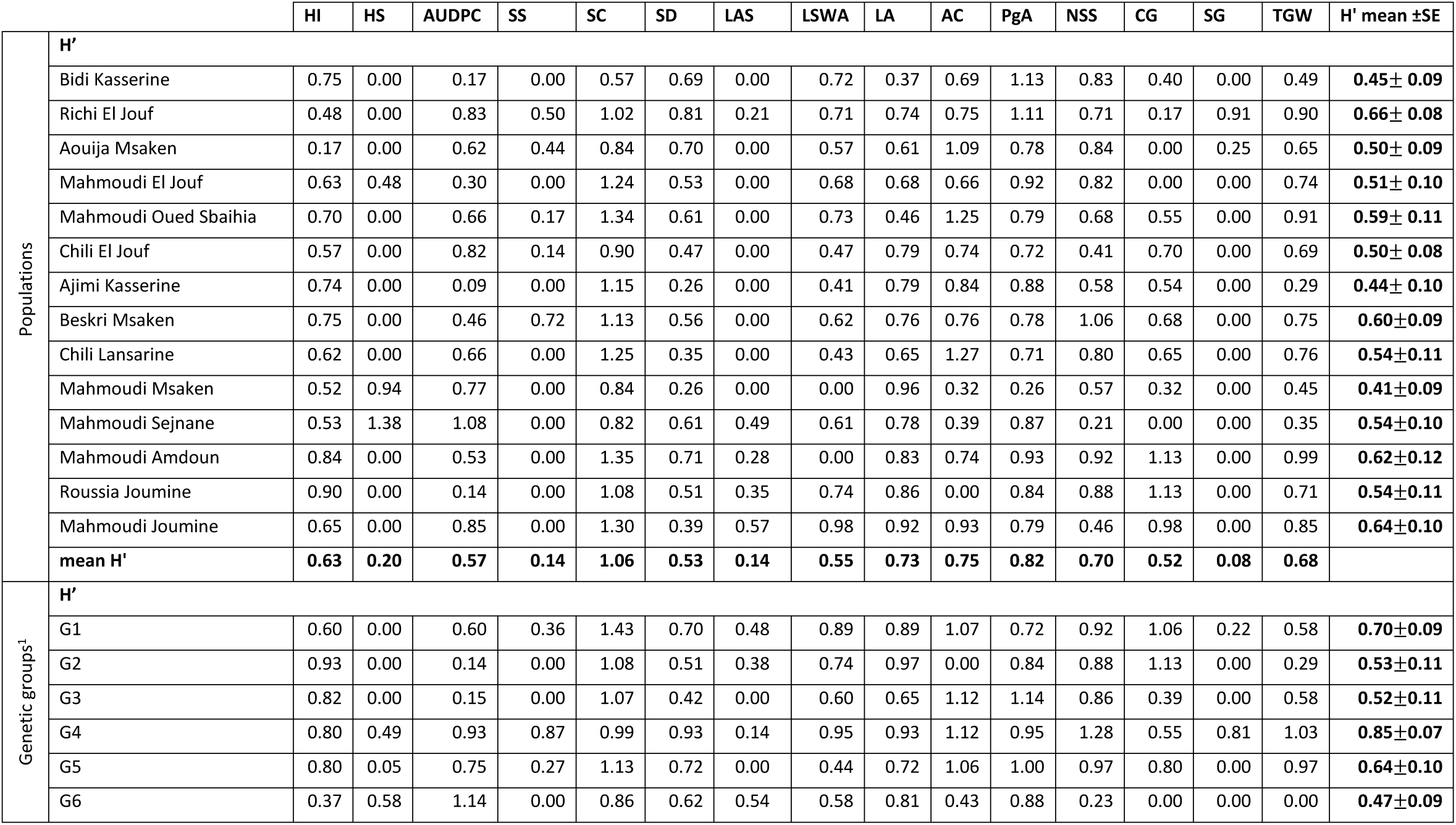
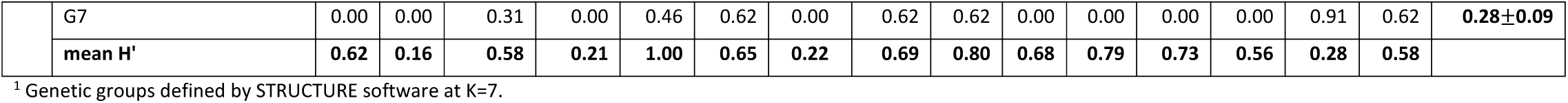
Estimates of Shannon–Weaver Diversity Index H’, mean H’ and standard error (±SE) of each of the 14 populations and seven genetic groups^1^ based on the evaluation of lineages with 15 phenotypic traits.

At the genetic group level, the greatest diversity (H’=0.85) was found in genetic group 4 composed by individuals belonging to Richi El Jouf, Mahmoudi Msaken and Aouija Msaken. Six from the 15 evaluated traits were highly polymorphic in this genetic group compared to others: SS (H’=0.87), spike density (SD) (H’=0.93), length of spike without awns (LSWA) (H’=0.95), awn colour (AC) (H’=1.12), number of spikelets per spike (NSS) (H’=1.28) and thousand grains weight (TGW) (H’=1.03). The genetic group 7 was the least phenotypically diversified (H’=0.28) but this group consists of only two lineages and is therefore not representative. The phenotypic diversity observed within the genetic groups indicate that they are not phenotypically homogeneous. Nevertheless, some phenotypic traits are clearly distinct between them. For example, SS with parallel-edge spikes being dominant in genetic groups 2, 3 and 7 (monomorphic H’=0) while less frequent in groups 1, 4 and 5. Genetic group 6 was characterized by a particular stunted-spike shape, which is not listed in the official classification from UPOV. Few individuals from genetic group 4 also had this particular stunted-spike shape. Moreover, SC was the most diverse phenotypic trait among all the genetic groups (mean H’ = 1.00).

#### -Statistics on quantitative traits

Nine phenotypic traits were quantitatively evaluated and subjected to statistical analyses. Significant positive correlations (at *p*<0.01) were detected between heading date (HS) and plant height (HI) (0.71), and not surprisingly between length of awns (LA) and length of awns in relation to spike (LAS) (0.82) (Figure-ESM2). Concerning *Septoria tritici* blotch (STB), a significant negative correlation (at *p*<0.01) was detected between area under disease progress curve (AUDPC) and HI (−0.43), and between AUDPC and HS (−0.57), corroborating that HI and HS can influence disease development. Early heading and short lineages tend to be more infected by STB. Finally, LSWA and SD were also significantly negatively correlated (−0.50).

Phenotypes of all quantitative variables were not normally distributed. Only the trait NSS could be normalized following a Box-Cox transformation (Box & Cox, 1964). A significant difference between population means was found for 8 quantitative traits (at *p*<0.001); LAS was not significantly different between population means. We observed a maximum of pairwise significant differences between populations for the trait NSS (35 pairs) and a minimum for the trait LA (five pairs) (Table 6).

**Table 6.**
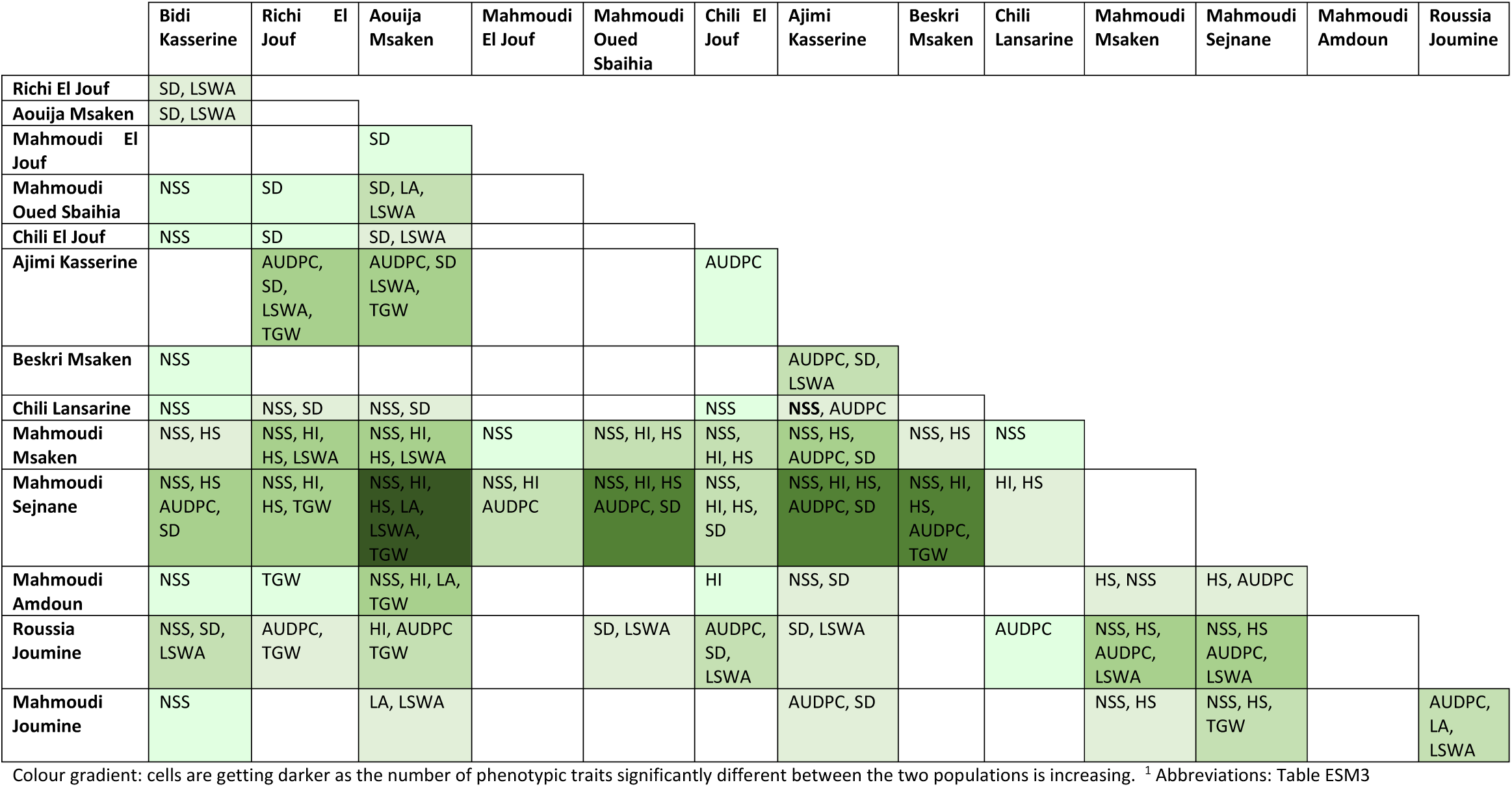
Pairwise populations significantly different quantitative traits^1^ at *p*<0.001.

The population Mahmoudi Sejnane cumulated the highest number of differences with other populations for quantitative phenotypic traits, except with the population Mahmoudi Msaken (Table 6). In total, 31 pairs of populations were not significantly different for any of the phenotypic traits studied, including the phylogenetically related (Figure 4) Richi El Jouf and Aouija Msaken, Mahmoudi Joumine and Beskri Msaken, Mahmoudi El Jouf and Mahmoudi Oued Sbaihia, Bidi Kasserine and Ajimi Kasserine, and even Mahmoudi Sejnane and Mahmoudi Msaken (Table 6).

#### -Response of the 14 populations to STB infection

The 14 durum wheat populations were contrasted in their response to STB infection (Figure 5). The two populations Ajimi Kasserine and Roussia Joumine show a nearly complete and qualitative resistance level to the *Z. tritici* strain IPO91009, while Mahmoudi Sejnane appeared highly susceptible. The other 11 populations show varying degrees of resistance or susceptibility to STB. Within each population, a variation in response to STB was also observed between lineages.

**Figure 5.**
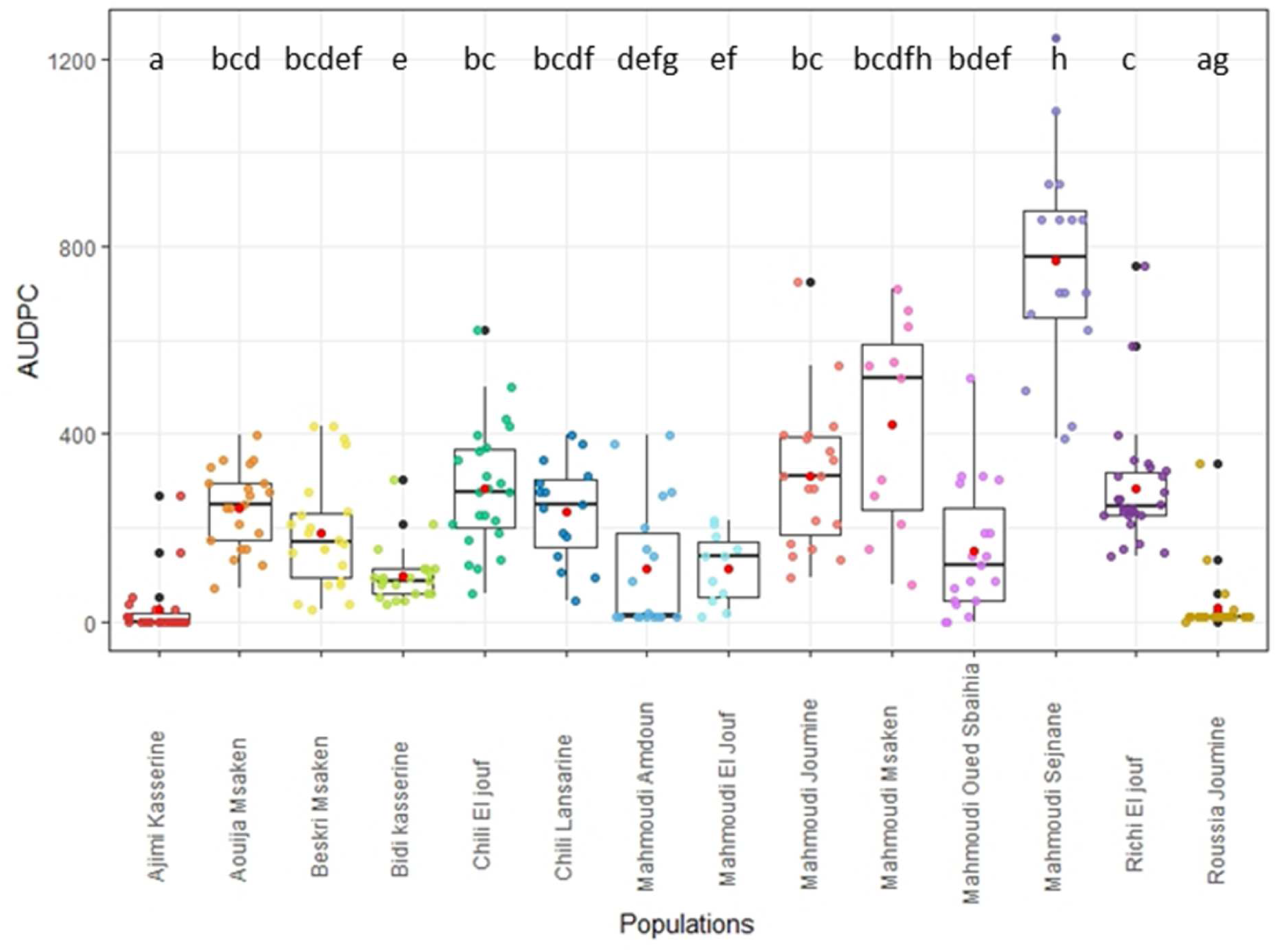
*Septoria tritici* blotch (STB) AUDPC (area under the disease progress curve) for the 14 durum wheat populations obtained after their inoculation in field conditions at Kodia Bou Salem with the *Z. tritici* strain IPO91009. Means are represented by a red point. Populations with significantly different means are indicated by different letters after Kruskal-Wallis and Mann–Whitney tests at *p*=0.05.

#### -Factor Analysis of Mixed Data (FAMD)

A Factor Analysis of Mixed Data (FAMD) was performed on both quantitative and qualitative variables (Figure 6). The first two principal components in the FAMD accounted for 16.5% and 9.8 % of the total variation, respectively, and together explained 26.3% of the total variation (Figure 6A). SS, HI, HS, NSS, TGW, AUDPC and LA were the most important traits contributing to the first principal component (Figure ESM3). Shape of grain (SG), SS, SD, AUDPC, TGW and LSWA contributed significantly to the second principal component (Figure ESM3). FAMD allowed to discriminate between three groups of populations. The first group includes Mahmoudi Sejnane and Mahmoudi Msaken (top left quadrant of Figure 6A), which are differentiated from the others populations by their stunted spike shape (Figure 6B), earlier heading date, shorter plant height, higher susceptibility to STB, and lower number of spikelets per spike (*p* <0.05). The second group includes Aouija Msaken and Richi El Jouf (top right quadrant of Figure 6A), which are distinguishable by their fusiform spike shape (Figure 6C), elongated shape of grains (Figure 6D), and the highest TGW compared to other populations (*p* <0.05). Finally, a third group includes the other 10 populations, which are characterized by spikes with parallel edges (Figure 6F) and semi-elongated grains (Figure 6E).

**Figure 6.**
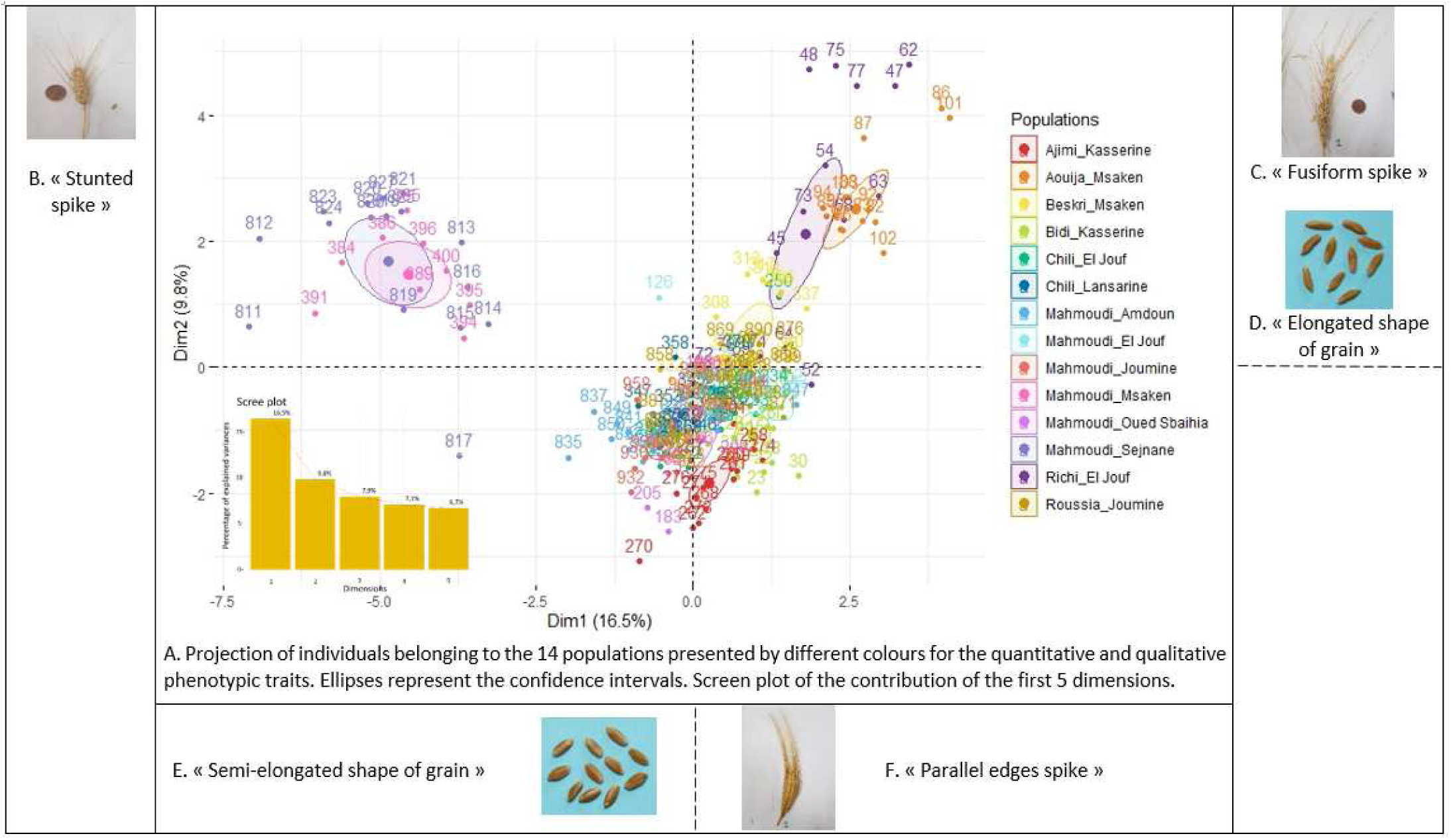
A: Factor Analysis of Mixed Data (FAMD) of 15 quantitative and qualitative phenotypic traits measured in the 14 durum wheat populations. B-F: Pictures of corresponding spike and grain shapes.

#### -P_st_-F_st_ comparaison and sensitive analysis

Comparisons of P_st_ and F_st_ show that most phenotypic traits are highly divergent reflecting therefore a local adaptation rather than a genetic drift, because lower 95% CI for P_st_ were higher than the F_st_=0.57 (Figure 7). Only for LAS, P_st_ didn’t differ from F_st_ since the lower 95% CI=0.5066<F_st_. However, the observed degree of differentiation is unlikely due to genetic drift for this trait (Leinonen & al., 2013). Evidence for the robustness of the sensitive P_st_-F_st_ analysis varied among the eight highly divergent phenotypic traits but was exceptionally strong for some traits such as HI, HS and AUDPC, which had critical c/h^2^ around 0.1 (Table 7). Similarly, LSWA, NSS and TGW have a critical c/h^2^ ranging from 0.16 to 0.21. Therefore, assuming that these traits are under selection and not under genetic drift is a very robust inference since phenotypic variance across populations that is explained by additive genetic effects would need to be at least six times lower than the phenotypic variation within populations to be explained by genetic drift. For SD and LA, the higher critical c/h^2^ of 0.30 and 0.61, indicates that phenotypic differentiation is lower and the conclusion of selection is clearly not conservative.

**Table 7.** P_st_–F_st_ comparison for quantitative traits measured in 273 lineages belonging to the 14 durum wheat populations. Lower values of the critical c/h^2^ indicate a more robust inference of local adaptation.

| Traits<sup>μ</sup> | P<sub>st</sub> at c/h<sup>2</sup> =1 <sup>§</sup><br>(null assumption) | F<sub>st</sub> | 95% CI lower | 95% CI upper | Critical c/ h<sup>2</sup> <sup>£</sup> |
| --- | --- | --- | --- | --- | --- |
|  |  | 0.57098 | 0.5218 | 0.6180 |  |
| HI | 0.9580 |  | 0.9353 | 0.9760 | 0.1118 |
| HS | 0.9692 |  | 0.9417 | 0.9881 | 0.1001 |
| AUDPC | 0.9506 |  | 0.9319 | 0.9680 | 0.1182 |
| SD | 0.9073 |  | 0.8432 | 0.9625 | 0.3008 |
| LAS | 0.5784 |  | 0.5066 | 0.7535 |  |
| LSWA | 0.9123 |  | 0.8904 | 0.9399 | 0.1991 |
| LA | 0.7732 |  | 0.7252 | 0.8585 | 0.6129 |
| NSS | 0.9299 |  | 0.9064 | 0.9537 | 0.1670 |
| TGW | 0.9220 |  | 0.8813 | 0.9551 | 0.2179 |

<sup>μ</sup> Abbreviations: Table ESM3 <sup>§</sup> c/h<sup>2</sup> =1 is the null assumption meaning that the proportion of phenotypic variance caused by additive genetic effects is the same for between-population and within-population variances (Brommer, 2011) <sup>£</sup> critical c/h<sup>2</sup> ratio was calculated according to the formula (Brommer, 2011).

**Figure 7.**
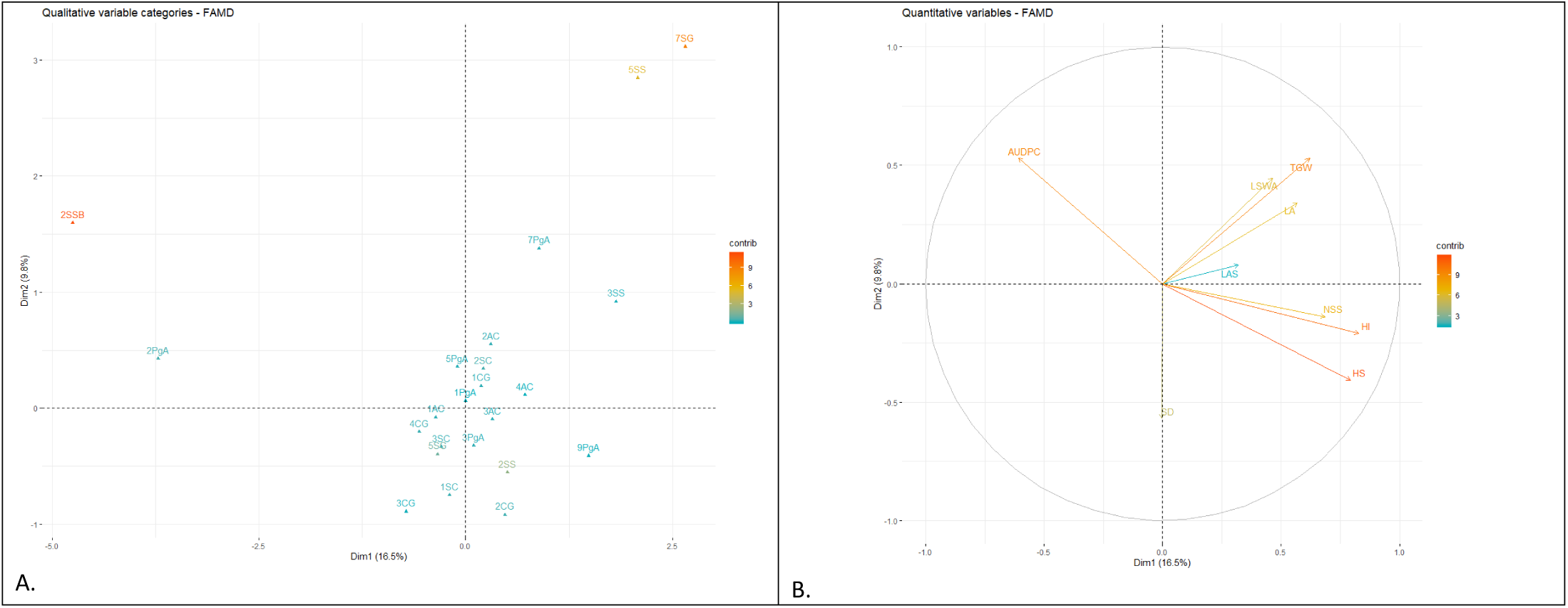
Output of the Factor Analysis of Mixed Data (FAMD). A. Categorical variable factor map projects the classes of qualitative variables in the plane of 1 and 2 dimensions underlining their contribution. Classes of Spike Shape: 1SS, 2SS, 2SSB, 3SS, 5SS; classes of Spike Colour: 1SC, 2SC, 3SC; classes of Awn Colour: 1AC, 2AC, 3AC, 4AC; classes of Anthocyanin colouration of Awns: 1PgA, 3PgA, 5PgA, 7PgA, 9PgA; classes of Colour of Grain: 1CG, 2CG, 3CG, 4CG; classes of Shape of Grain: 1SS, 2SS, 3SS. **B. Correlation circle represents the projection of quantitative variables on the 1 and 2 dimensions underlining their contribution**. HI: plant height; HS: heading date; AUDPC: area under disease progress curve; SD: spike density; LAS: length of awns in relation to spike; LSWA: length of spike without awns; LA: length of awns; NSS: number of spikelets per spike; TGW: thousand grains weight.

**Figure 8.**
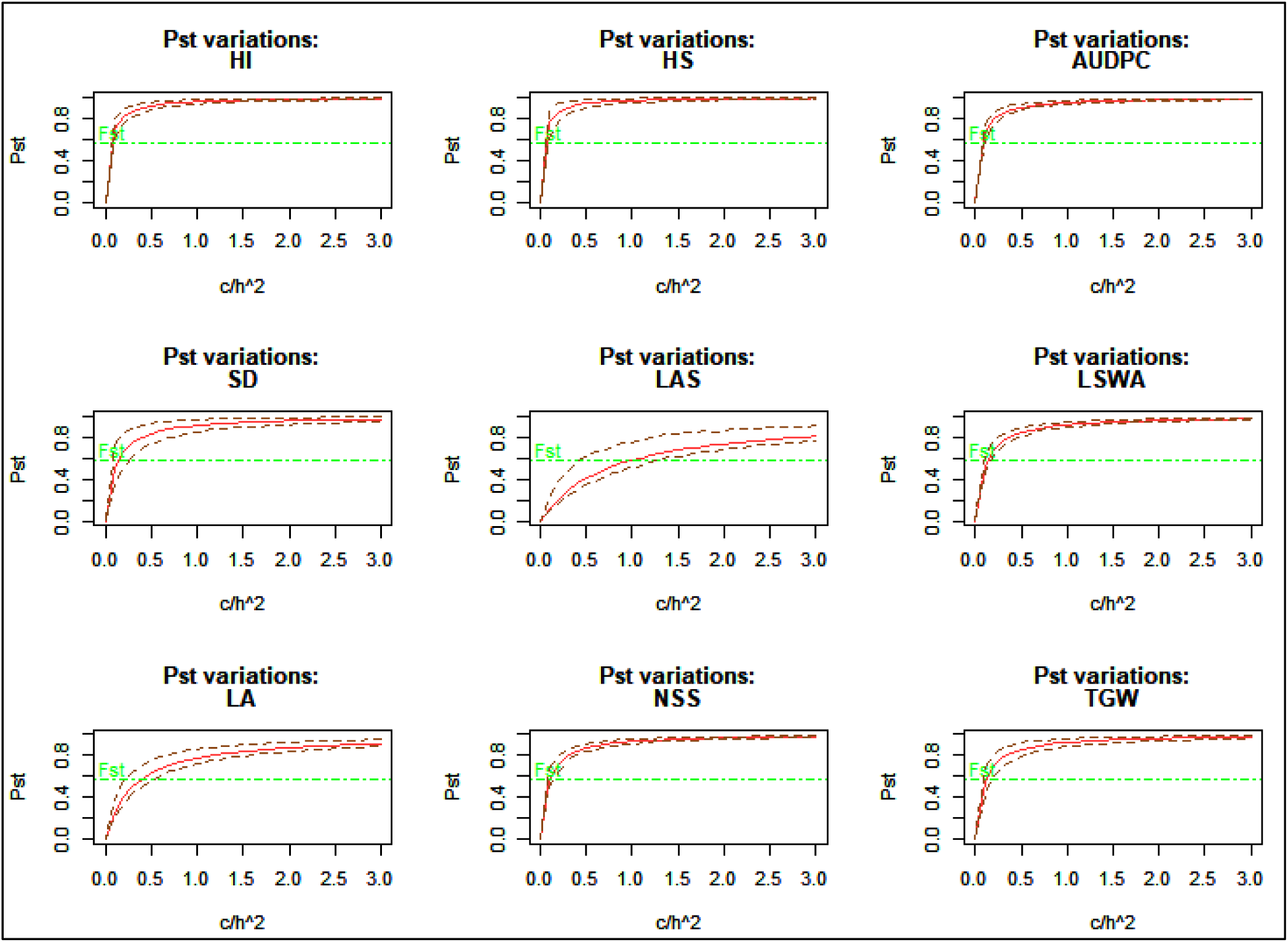
Comparison of phenotypic differentiation of the 14 durum wheat populations (P_st_: red solid line; dotted red lines represents the 95% CI). The neutral genetic differentiation (F_st_-dotted green line for F_st_=0.57098 was calculated using the package hierfstat with Rstudio version 3.5.2), while the ratio c/h^2^ ranged from 0 to 3 where c represents the proportion of the total variance and h^2^ the heritability. The lowest value of c/h^2^ for which P_st_ exceeds F_st_ = critical value of c/h^2^ can be considered an indication of the robustness of using P_st_ as an alternative for Q_st_ (Brommer, 2011).

## DISCUSSION

Durum wheat landraces traditionally used by Tunisian farmers have been progressively replaced by modern cultivars provided by international (i.e. CIMMYT) and Tunisian (i.e. INRAT) breeding programs. This led to a rapid increase in productivity of wheat and was accompanied by the multilateral trade liberalization on its value chain as well as staple food affordability in both rural and urban Tunisian areas (La Rovere & al., 2010). However, few farmers are still until now cultivating landraces in some mountainous or dry areas of Tunisia characterized by low input farming systems, for their own consumption or to be sold on local markets. Understanding the genetic and phenotypic diversity of landraces can be used to legitimate the preservation of these important genetic resources in order to improve the adaptability of durum wheat crop to marginal environments and threatening pathogens (Pietrusińska & al., 2018).

### 1- Durum wheat landraces are genetically diverse and dynamic populations

#### -Microsatellite markers

PIC values are a good indication of the informativeness of the multiplexed panel of the microsatellite (SSR) markers used in this study. According to the classification by Botstein & al. (1980), our nine SSRs were moderately (0.25<PIC<0.5) to highly (PIC>0.5) informative, therefore sufficient to discriminate between populations and useful for further genetic diversity studies and seed bank management.

The genetic diversity of the 14 populations assessed with these SSRs (He=0.57) was almost equivalent to the genetic diversity of several collections of durum wheat landraces from different Mediterranean countries such as Tunisia, Morocco or Italy with average gene diversity values between 0.44 and 0.69 (Nefzaoui & al., 2014; Slim & al., 2019; Sahri & al., 2014; Soriano & al., 2016). This high genetic diversity can explain why Tunisia has previously been considered as a centre of diversity for durum wheat (Ayed & al., 2010).

#### -Repartition of genetic diversity

The Analysis of MOlecular Variance (AMOVA) evidenced the great genetic diversity present within the durum wheat populations since variation was only slightly greater between populations (54%) than within population (45%) (Table 2). It can be the result of outcrossing and fitness-relevant mutations. The global F_st_ value of 0.5 also supports a high differentiation between the 14 populations. Similar results were obtained for Ethiopian durum wheat landraces with a slightly higher genetic variance between populations (52%) than between individuals within genetic groups (48%) (Alemu & al., 2020). However, several other studies rather report a greater genetic variance within than between populations (Kabbaj & al., 2017; Soriano & al., 2016; Kyratzis & al., 2019; Asmamaw & al., 2019), potentially explained by seed exchange or selection by farmers for similar traits.

#### -Structuration of lineages

Based on a bayesian method, the 14 populations were stratified into seven genetic groups (Figure 2). Only few lineages showed a low membership coefficient q-value suggesting a weak level of admixture among the populations. This is probably due to the fact that wheat is mostly autogamous and to the selection pressure exerted by farmers. Some genetic groups are composed of lineages belonging to a single population while other genetic groups include lineages from different populations. Two genetic groups (G2 and G6) are exclusive to a single population (Roussia Joumine and Mahmoudi Sejnane, respectively), as supported by the distant position of each population on the minimum spanning network (Figure 3). Roussia Joumine is the only population with variety name ‘Roussia’, and this landrace is likely to be genetically distinct from the others. Mahmoudi Sejnane seems unrelated to other ‘Mahmoudi’ populations. This is also true for Mahmoudi Msaken composed only from lineages belonging to a group (G4) which also includes individuals from the Richi El jouf and Aouija Msaken populations. Another group (G5) is composed of lineages coming from eight populations, including nearly all individuals from Chili El Jouf and Chili Lansarine. This suggest that both populations probably have a common origin and thus highlights the pronounced impact of the variety name in the genetic structuration. Furthermore, a group (G3) is composed of lineages coming from four populations, including nearly all individuals from Ajimi Kasserine and Bidi Kasserine. This strong evidence of the impact of locality on the genetic of Tunisian durum wheat landraces structuration, in addition to the expected effect of the variety name, has been demonstrated in previous studies (Slim & al., 2019; Soriano & al., 2016; Rufo & al., 2019). Finally, a group (G1) is composed of lineages coming from seven populations having different origins and different names of variety, underlining the more complex genetic structure of several populations. Genetic structuration of these 14 durum wheat populations is therefore explained by a combination of effects, including the locality of origin and the variety name, that likely results from the farmer-mediated selection and exchange trajectory of each population.

#### -Agro-morphological diversity

Characterizing the phenotypic diversity of durum wheat landraces, known to be often agro-morphologically heterogeneous (Royo & al., 2017), is prominent towards an understanding of their life history, in other words the selection trajectory and seed exchanges between farmers. Indeed, the 14 studied populations were agro-morphologically diversified with a mean Shannon-Wiener index of H’=0.54, estimated with 15 phenotypic traits. In previous studies, even greater agro-morphological diversities were observed with H’=0.77 within 30 Tunisian durum wheat landraces characterized for 11 phenotypic traits (Ayed & al., 2010) and H’=0.63 within a collection of 368 Tunisian durum wheat accessions characterized for 9 phenotypic traits (Slim & al. 2011). The least diversified traits among the 14 populations were SG (shape of grain) (H’=0.08), SS (spike shape) (H’=0.14) and LAS (length of awns in relation to spike) (H’=0.14). The most frequent shape of grains is semi-elongated, which is associated with a spike shape with parallel edges. Thus, it suggests a selection pressure exerted by Tunisian farmers on their landraces for the same spike shapes. However, grain and spike shapes appeared highly diversified in several other studies (Belhadj & al., 2015; Bechere & al., 1996; Al Khanjari & al., 2008). The most diversified traits among the 14 populations were SC (spike colour) (H’=1.06), PgA (anthocyanin colouration of awns) (H’=0.82) and AC (awn colour) (H’=0.75) (Table 6). Spike and awn colours are important for Tunisian farmers. For example, farmers associate the black colour of awns with the Mahmoudi landrace distinguished for its higher quality for semolina and bread bakery. These findings are consistent with previous studies showing that awn colour was often among the most diversified traits in Tunisian and Omani landraces (Belhadj & al., 2015; Al Khanjari & al., 2008).

Population genetic groups did not appear congruent with the structure of agro-morphological diversity while another study highlighted a reliable relationship between genetic and phenotypic population structures, and the connection of both with the geographic origin of the landraces (Soriano & al., 2016). The overall agro-morphological diversity measured with Shannon index does not differ significantly between the 14 populations. A better discrimination between the populations could be obtained with more informative traits and tighter descriptors. Traits related to spike and awn colours could be described using a colour palette for better discrimination between shades, such as Roussia Joumine’s russet colour of awns. A broader list of spike shapes could be established including for example the stunted form of spikes found in Mahmoudi Msaken and Mahmoudi Sejnane populations.

### 2- Assessing the origin of durum wheat landraces

Characterization of the 14 populations revealed the complexity of the history of durum wheat cultivation in Tunisia. Behind each population hides a particular life story that can only be understood in light of converging evidences from genetic and agro-morphological descriptions of the populations, and from their local context.

#### -Roussia Joumine, standing-out redhead

The population Roussia Joumine, the only one with variety name ‘Roussia’, stood-out for the homogeneous russet colour of spikes from all its characterized lineages, although this criterion is not in the UPOV classification. It was also strongly genetically distinguishable from all the other studied populations. In literature, Roussia 875 is a local landrace from the north of Tunisia identified in 1927 and registered from 1953 to 1973 (Ammar & al., 2011). Gharbi & El Felah (2013) reported that ‘Roussia’, keep its name from an original population that was cultivated extensively in field crops. Slim & al. (2019) previously reported a dendrogram for the durum wheat collection of the NGBT on which Roussia Joumine and Mahmoudi Amdoun clustered together, while in our study both populations were highly differentiated (F_st_=0.6). Indeed, Roussia Joumine showed higher values for all genetic indices evaluated, and it was the most genetically diversified population while being phenotypically less divergent. This landrace was as well remarkably resistant to STB and could be an interesting source of resistances to this disease.

#### -Chili El Jouf and Chili Lansarine, a common origin

The two populations Chili El Jouf and Chili Lansarine, were significantly weakly differentiated with SSR markers (F_st_=0.03) and almost exclusively composed of lineages classified in the same genetic group (Figure 2) and sharing an MLG (Figure 3). Both populations were grouped with ‘Chili’ landraces collected by USDA in Tunisia in 1979 (Figure 4). In literature, the origin of ‘Chili’ landrace is not very clear. According to Saade & al (1996) ‘Chili’ was selected from a commercial shipment imported from Chile in the early 1960s and rapidly adopted by Tunisian farmers. Deghais & al. (2007) reported that, in 1932, sir Racine, a French industrialist, sent from Marseille (France) 100 kg of Chili seeds to sir Charles Fabre, a farmer in Bou Salem. In 1938, the Botanical Service of Tunis recovered a part of these Chili seeds and incorporated them in regional field trials to evaluate its adaptative potential to regional conditions. Propagation of Chili then took place as early as 1941/1942 and the Botanical Service of Tunis selected the variety ‘Chili-931’, which was registered in the official Tunisian catalogue of varieties in 1953. Since 1985, the cultivation of this variety has been largely abandoned with the exception of a few niches and it was removed from the catalogue in 1993.

Chili El Jouf and Chili Lansarine were identical for most phenotypic traits studied, except for the NSS, being higher in Chili El Jouf than in Chili Lansarine (Table 6). The difference between the two populations could result from a selection by the farmers from El Jouf for a better yield, whose NSS is a component. Moreover, as NSS is influenced by environmental conditions, such as temperature and day length (Knezevic & al., 2007; Rawson, 1970), the difference between the two Chili populations could also have resulted from a local adaptation to environmental conditions. Genetic analysis demonstrated that Chili El Jouf had lower genetic diversity than all studied populations (Table 3).

#### -Aouija Msaken and Richi El Jouf, distant cousins

The populations Aouija Msaken and Richi El Jouf stood out for their similar fusiform spike shape and elongated grain shape (Figure 6). These populations are genetically heterogeneous and share lineages belonging to the same genetic groups (Figure 2) but remain genetically differentiated from each other (F_st_=0.27). Based on Neighbour joining clustering, Richi El Jouf and Aouija Msaken were genetically closely related (bootstrap support of 85.2), with an F_st_ index around 0.2 being the lowest between Richi El Jouf and any other population. In addition, both populations share a common MLG (Figure 3) suggesting that they have a common ancestor. Richi is a very old variety related to the species *Triticum polonicum* (personal communication from NGBT). According to some studies, Aouija is also called “Aouej”, “Swabaa Aljia” or “Aouiji”. Several forms were introduced in 1913 and 1916 by Boeuf from Morocco where they were called “Zréa Laouaja” in relation to the hunchback shape of the grain reminiscent of that of *Triticum polonicum*. Richi El Jouf was the most diversified population based on all measured traits (H’=0.66), while Aouija Msaken had a lower phenotypic diversity (H’=0.50).

#### -Ajimi Kasserine and Bidi Kasserine, from the same bag

The two populations collected in Kasserine, Bidi Kasserine and Ajimi Kasserine, were undifferentiated from each other, genetically (F_st_=0.039) and phenotypically for quantitative traits. Both of them shared three MLGs, had exclusively the same heading dates, same parallel edged spikes and same spike shapes. Some lineages from Ajimi Kasserine were shorter, with shorter and more compact spikes, longer awns and lower number of spikelets per spike, albeit statistically not significant. In addition, some lineages of Ajimi Kasserine had white colour of seeds and spikes and high anthocyanin pigmentation of awns that was not observed in Bidi Kasserine. The populations Ajimi Kasserine and Bidi-Kasserine were genetically close based on neighbour joining clustering (bootstrap support of 100, Figure 4) and on their F_st_ index around 0.04. These results support the hypothesis of a common origin for these two populations, considering that the NGBT collected seed samples from these two populations from the same farmer in the region of Kasserine. Our findings are consistent with an interview with the farmer suggesting a mishandling of seed bags that led to name two bags with different variety names while they originated from the same landrace.

#### -Majmoudi Sejnane, a fake identity

Mahmoudi Sejnane was highly differentiated from all the other ‘Mahmoudi’ populations (F_st_: 0.53-0.8), but was phenotypically identical to Mahmoudi Msaken. Both, Mahmoudi Sejnane and Mahmoudi Msaken, were genetically homogeneous with lineages belonging to only one genetic group each, respectively G4 and G6. These two populations were characterized by their stunted spike shape and higher susceptibility to STB. The history of Mahmoudi landraces is unclear and several landraces may have been wrongly attributed the variety name ‘Mahmoudi’ by farmers. In fact, it appears that the population Mahmoudi Sejnane is genetically very close from the modern variety ‘Karim’ (bootstrap 88.4, Figure 4). A recent study by Slim & al. (2019), studying the same population Mahmoudi Sejnane and the modern variety ‘Karim’ by neighbour joining cluster analysis, reached the same conclusion. Thus, the population Mahmoudi Sejnane is suspected to be derived from the modern variety ‘Karim’. The short plant height of Mahmoudi Sejnane (and Mahmoudi Msaken) compared to other populations and its high level of susceptibility to STB strongly looks alike the characteristics of the variety ‘Karim’, which has been introduced in Tunisia more than 40 years ago by CYMMIT. It is by far the most cultivated variety in Tunisia. Its resistance to STB has undergone years of genetic erosion making it today very susceptible to *Z. tritici*.

#### -Mahmoudi Msaken, a probable ‘Chili’ origin

Mahmoudi Msaken was highly differentiated from all the other populations with variety name ‘Mahmoudi’. The phylogenetic tree revealed that Mahmoudi Msaken is rather strongly related to a landrace carrying the name ‘Chili’ (bootstrap support of 98.9), which do not group with other ‘Chili’ landraces (Figure 4). However, the short length, stunted spike shape (Figure 6) and higher susceptibility to STB looked alike the characteristics of Mahmoudi Sejnane, while these two populations were genetically highly differentiated (F_st_=0.8). This suggests that Mahmoudi Msaken is inherited from the landrace Chili (PI534336, 1976) but might have evolved through mutations and crosses with other varieties, such as the widespread variety Karim. This hypothesis could be investigated further through the phenotypic characterization of the historical landrace.

#### -Beskri Msaken, a ‘Mahmoudi’ on disguise

Beskri Msaken was composed of lineages belonging to the same genetic groups (i.e. G1 and G5) as other populations with the variety name ‘Mahmoudi’ (Figure 2). Moreover, Beskri Msaken was weakly differentiated from the populations Mahmoudi El Jouf and Mahmoudi Joumine, these populations sharing two MLGs. The close relationship between ‘Beskri’ and ‘Mahmoudi’ landraces was previously established (Robbana & al., 2019; Slim & al., 2019). Based on Neighbour joining clustering, the populations Beskri Msaken and Mahmoudi Joumine also appeared genetically close, even when the F_st_ index was around 0.5. It suggests a common ancestor between Beskri Msaken and other Mahmoudi populations.

#### -’Mahmoudi’, a popular variety with a complex history

The four populations Mahmoudi El Jouf, Mahmoudi Oued Sbaihia, Mahmoudi Joumine and Mahmoudi Amdoun had a heterogeneous genetic structure but all have lineages belonging to G1 and G5 (Figure 2). In addition, Mahmoudi Oued Sbaihia and Mahmoudi Joumine had lineages belonging to G3. Mahmoudi Amdoun was genetically differentiated from the three other populations with variety name ‘Mahmoudi’ (F_st_ 0.247-0.346) but remained identical phenotypically. The genetic differentiation between ‘Mahmoudi’ populations from El Jouf and Oued Sbaihia, and from El Jouf and Joumine, was weak while they did differ phenotypically for quantitative traits. The complexity of the genetic structure of ‘Mahmoudi’ populations emphasises the complexity to decipher and identify the determinants of the life history, *i*.*e*. selection trajectory and seed exchanges between farmers, of those landraces. In the literature, ‘Mahmoudi’ landraces were reported from Tunisia, Algeria, Libya or Palestine (Deghais & al., 2007; Erroux, 1991; Boeuf, 1926; Gharbi & El Felah, 2013; Ammar & al., 2011). In Tunisia, Boeuf (1926) reported seven indigenous botanical Mahmoudi varieties that were selected and propagated by the Botanical Service of Tunis in the early 1900s. In the middle of the twentieth century ‘Mahmoudi’ landraces became the most commonly cultivated landraces by traditional farmers in Tunisia, even exclusively in certain locality such as Joumine. Farmers questioned during a survey appeared very attached to this landrace for its yield stability, and high quality for making couscous and traditional bread. Based on pairwise F_st_ analysis, Chili (PI534351, 1976) was almost genetically identical to Mahmoudi Joumine (F_st_=0), Mahmoudi El Jouf (F_st_=0) and Mahmoudi Oued Sbaihia (F_st_=0.04). These three varieties named ‘Mahmoudi’ were also weakly differentiated from Chili Lansarine and Chili El Jouf, suggesting that the landrace ‘Chili’ has contributed to the complex structure of ‘Mahmoudi’ landraces in Tunisia. On the phylogenetic tree with historical collections of Tunisian landraces, these three varieties named ‘Mahmoudi’ populations and Mahmoudi Amdoun, clustered with landraces carrying the variety name ‘Chili’ rather than with landraces carrying the variety name ‘Mahmoudi’ (Figure 4). It strengthens the hypothesis that seeds from ‘Chili’ have been introduced into ‘Mahmoudi’ landraces. Also, the success of ‘Mahmoudi’ in Tunisia may explain why many seed lots (or populations) have been called by traditional farmers with the name ‘Mahmoudi’ while having different origins, resulting therefore in the complex genetic and phenotypic structure that we observed. Overall, our findings emphasise the fact that the name of varieties is not sufficient to explain the origin and history of Tunisian durum wheat landraces.

### 3- Challenges and opportunities in the conservation of durum wheat landraces

#### -Contamination of durum wheat landraces with a hexaploid wheat

Our study revealed the presence of hexaploid lineages in eight out of the 14 studied populations, easily distinguishable in the farmers’ fields because of their long cylindrical white spikes. They are considered as impurities and result in a semolina with a flour-like texture with poorer quality. The origin of this species in Tunisian durum wheat fields is not well understood. Some authors describe it as indigenous and others as introduced by Europeans (Bœuf, 1926; Portères, 1958), while more recent studies established that durum wheat fields grown with landraces are frequently contaminated with hexaploid wheats (Zeven & Waninge, 1989; Figliuolo & al., 2007). In the middle of the twentieth century the proportion of bread wheat in Tunisian durum wheat fields approached 50% (Bœuf, 1926; Portères, 1958), but this proportion is lower nowadays and was only 9% in our study. These bread wheats or ‘mule’s tail*’* existing as a mixture in the durum wheat fields are locally called " Baabous Bghal " = " *Babous el brel "*, " Dhil Bghal " or " Boujlida " = *" Bou jelida* " (in English: mule’s tail), and Bœuf and collaborators were unsuccessful in domesticating them through breeding (Bœuf, 1926; Gharbi & Elfalah, 2013). Portères (1958) suggested that this form already existed before the arrival of Europeans, and that durum wheat brought by Arabs in North Africa competed bread wheat due to its resistance to drought and its better use for semolina. He further mentioned that the Romans, perhaps the Phoenicians and Greeks, brought bread wheat to North Africa. Ducellier described in 1923 an Algerian wheat named ‘Hachadi’ (white, dense, very rough ear; strongly curved, inflated glumes; strongly spreading ears; strong divergent beards; reddish or reddish-brown seeds), which might be synonymous to Tunisian ‘mule’s tail’ (Laumont & Erroux, 1962). ‘Hachadi’ is probably an old form of cultivated wheat, which is nowadays never found in its pure state but always mixed with traditional durum or bread wheats. Nevertheless, these ancient forms have no cultural value anymore and are not interesting for breeders. Similarly, Laumont & Erroux (1962) reported the presence of a hexaploid wheat form in Algerian durum wheat fields, known as ‘Guelia’ (root akli: to cook, because of the reddish colour of the grain which would have been scorched in the heat). This form of wheat has spread in Algeria since 1950 as a result of the importation of wheat (Florence × Aurore) by Tunisian flour mills. It looks like the " Baabous Bghal " or " Boujlida " (Laumont & Erroux, 1962), is morphologically different from ‘Hachadi’ and characterised by rough, white ears, with a keeled husk similar to durum wheat, and with a red grain often confused with durum wheat grains.

The presence of these undesirable hexaploid wheats in durum wheat fields obliges Tunisian farmers to conduct a dynamic selection in order to eradicate ‘mule’s tail’, unsuccessfully so far. Farmers who are not undergoing this selection process are gradually losing the purity of their landrace, which can lead to a loss of its diversity and sometimes to its complete loss due to resignation.

#### -Divergence among landraces is explained by selection rather than genetic drift

The majority of evaluated quantitative traits were highly divergent (P_st_>F_st_), with a robust approximation of P_st_. Especially, HI, HS, AUDPC, NSS and LSWA were distinguishable with low critical c/h^2^ < 0.20. Indeed, we can suppose that divergence in genes coding for these five traits exceeds what is expected on the basis of genetic drift. It indicates that selection has resulted in different phenotypes for these five traits between the studied populations, which is the footprint of local adaptation. Selection made by farmers on their landraces should have contributed to this local adaptation. Indeed, a local survey informed us that some farmers perform irregular selection based especially on spike length, spike density and awn colour. Thus, the genetic variation present in the landraces for these traits constitute an important resource to breed for improved varieties adapted to local constraints and meeting Tunisian farmers’ expectations.

#### -Use of most resistant lineages to STB for conservation and breeding purposes

Landraces are a well-known source of resistance to diseases (Akem & al. 2000; Nazco & al., 2012; Xu & al., 2018; Agnoun & al., 2019; Yao & al., 2019), such as durum wheat landraces and their resistance to *Septoria tritici* blotch (STB) caused by *Z. tritici*. Previous studies by Ferjaoui & al. (2011, 2015) identified for instance new resistance genes to STB in ‘Azizi27’, ‘Agili37’, ‘Agili39’ and ‘Derbessi12’ landraces. Ouaja & al. (2020) also identified valuable sources of resistance among a collection of 304 Tunisian durum wheat landraces. In our study, the 14 populations responded differently to STB (Figure 5). Roussia Joumine and Ajimi Kasserine were the most resistant populations to the *Z*.*tritici* strain used for inoculations, IPO91009. The resistance is qualitative (total resistance controlled by major genes) in both populations suggesting the presence of highly effective resistance genes that could be useful for breeding. Bidi Kasserine appeared slightly less resistant to STB than Ajimi Kasserine, although the two populations are genetically and phenotypically identical. On the contrary Mahmoudi Sejnane was highly susceptible to STB, suggesting that it does not carry any source of resistance to this disease or that the resistance genes it may contain have been overcome by *Z. tritici* populations. Intermediary, the remaining 10 populations expressed varying levels of resistance, probably quantitative (partial resistance controlled by polygenes with moderate-to-small effects).

Intra-population variation of the level of resistance between lineages was observed as well (Figure 5), reflecting the genetic diversity previously described within the populations. Few lineages from Roussia Joumine and Ajimi Kasserine had some level of susceptibility. Inversely, several populations expressing quantitative resistance contained few completely resistant lineages, such as Mahmoudi Amdoun and Mahmoudi Oued Sbaihia. Moreover, all populations expressing quantitative resistance as a whole were composed of lineages with varying levels of resistance, which was especially pronounced for Mahmoudi Msaken. This intra-population diversity represents as well an important resource of genetic variation, which should not be overseen.

## Conclusion

Our study showed a complex structuration of 14 durum wheat populations, only partly explained by their geographic origin and name of variety. Among them, landraces called ‘Chili’ contributed a lot to the history of Tunisian durum wheat landraces. These results highlight that a landrace is the outcome of a complex selection trajectory, driven not only by the heritage of a biological material (seed lots) but also by traditional breeding practices (selection) deeply-rooted in a territory and subject to several kinds of disruptions (exchange, mixing, loss, ingression of exogeneous material, misidentification, etc.). These two components are an integral part of the “life history” of landraces. The dynamic nature of this life story must be considered and characterized, even more thoroughly than we have done in this study, for instance with sociologists, if one wants to better exploit the resources that landraces represent. It is also crucial to enlarge prospection and collection efforts of actual cultivated landraces to retrace the history of these genetic resources and, particularly, distinguishing the “real” landraces from the “false” by combining genetic and phenotypic approaches. The conservation of durum wheat landraces is important for future breeding programs. And the role of gene banks is prominent to develop appropriate and relevant *in situ* and *ex situ* conservation plans.

## Materials and Methods

### Plant material

Since 2012 the National Gene Bank of Tunisia (NGBT) conducts a program aiming at the conservation of landraces from traditional Tunisian crops, such as durum wheat. Farmers still cultivating those landraces were identified and seed samples were collected for conservation purposes. Within this framework, we focused on 14 populations, 10 being collected in 2015 from five localities and four being collected in 2017 from three additional localities (Table 8). Each population was considered as representative of a given durum wheat landrace and was designated by a variety name combined with its locality of origin. Therefore, several of these “variety-locality” populations collected in different “localities” have the same “variety” name, and vice-versa. Seeds of each population were randomly selected from the seed samples and sown under open field conditions in micro-plots (2 m × 1 m) at the experimental station of Thiverval-Grignon, France (48°50’21.3” N, 1°57’07.2” E). For each population, 32 plants were randomly selected and labelled from 1 to 32, except for the four populations collected in 2017 for which 16, 23, 38 and 51 plants were selected, leading to a total of 448 plants. At the inflorescence emergence stage (Zadoks’ 50-59), the wheat spikes were bagged in permeable cellophane wraps in cellophane to promote self-fertilization and prevent the expression of heterozygosity (< 5% in cultivated wheat). When matured (Zadoks’ 90-99), spikes from each plant were collected and stored in separate bags before being threshed, spike by spike. This procedure was repeated twice for the 10 populations collected in 2015 and only once for the four populations collected in 2017. As each cycle of self-fertilization is expected to reduce natural intercrossing by 50%, the residual heterozygosity was estimated less than 1.25%. Each lineage obtained from the 32 plants originally selected and represented by seeds coming from a single spike were sown as a single head-row during the growing season 2017-2018 on the CRP-Wheat Septoria Phenotyping Platform from CIMMYT-IRESA (Kodia Bou Salem, Tunisia). This trial was conducted as randomized incomplete block design (12 blocks of 16 m × 1 m separated by the modern variety ‘Karim’). Eight reference varieties were included in each block: ‘Karim’, ‘Khiar’, ‘Maali’, ‘Nasr’, ‘Salim’, ‘Agili-39’ and ‘Mahmoudi-101’. At the end of the trial, six spikes per wheat lineage (i.e. head-row) were harvested, threshed, and stored for further characterizations.

**Table 8.**
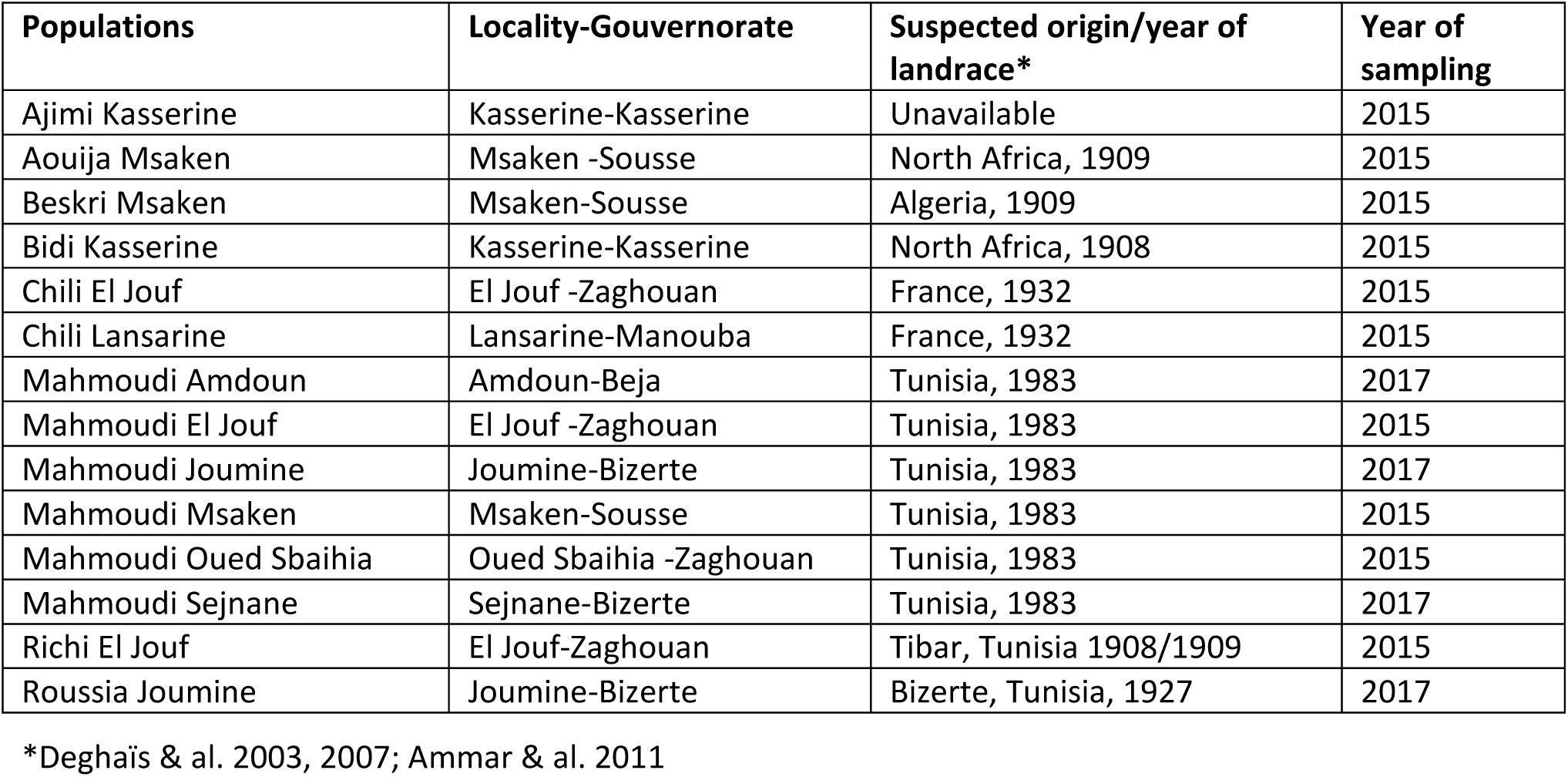
List of the 14 durum wheat populations: locality, suspected origin/year and year of sampling.

### Molecular analyses

#### *DNA isolation and genotyping

For each wheat lineage, one seed was sown and grown under controlled conditions (growth chamber) with a 16h/24h light (300 photons μmol.m^−2^.s^−1^) and a temperature of 18°C night/20°C day. Two weeks after sowing, a 2-3 cm segment from the youngest leaf of each plant was sampled for DNA extraction performed using the DNA plant MiniKit protocole (www.qiagen.com). Purity and concentration of the extracted DNAs were estimated using a Nanodrop spectrophotometer (ND-1000). The DNA samples were normalized to a concentration of 20 ng.μL^−1^ and genotyped using nine SSR markers (Table 9). Genotyping was outsourced at Eurofins (www.eurofins.fr). These SSR markers were assembled in a single multiplex following the methodology described by Gautier & al. (2014). Briefly, the forward primers were labelled with four fluorochromes (i.e. Ned, Fam, Pet, Vic) in a way that the same colour was given only to markers with non-overlapping range of allele sizes. Candidate markers were individually tested on eight reference durum wheat DNAs, before constitution and validation of the multiplex. Chromatograms were visually inspected for all markers and for all individuals using Peak scanner software version 1.0, before final assignment of SSR alleles. Individuals with missing data for at least one marker were removed from our dataset not to influence the detection of unique multilocus genotypes (MLGs) per population. Finally, genotypes of 335 lineages were obtained (13 to 32 lineages per population), after the exclusion of hexaploid lineages (see below, section “Determination of the ploidy level in lineages”).

**Table 9.**
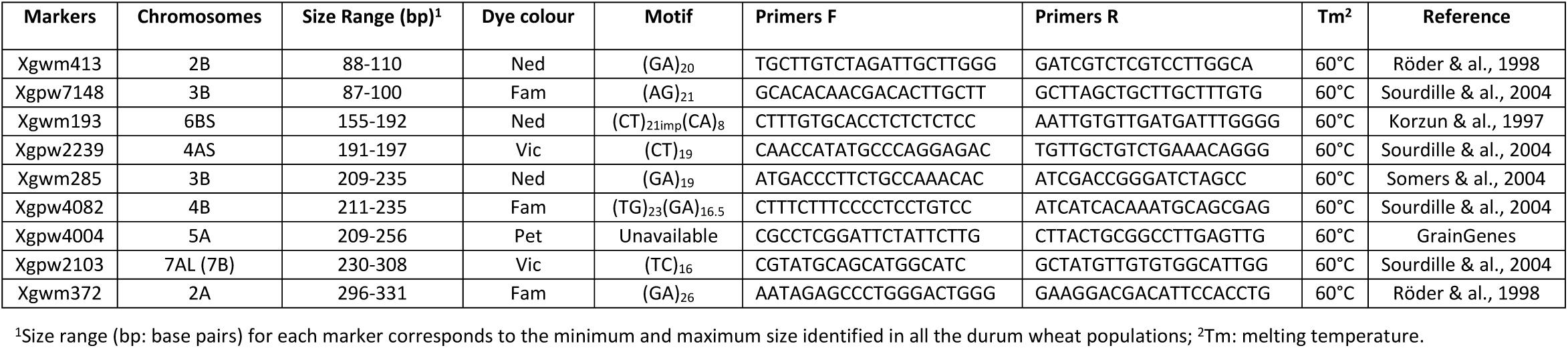
Description of the panel of 9 SSR (microsatellite) markers used for population genetics.

#### *Marker characterization

Neutrality of the SSR markers was preliminary confirmed using BayeScan v2.1 software, which allows identifying candidate loci under natural selection from genetic data using differences in allele frequencies between populations. The default settings were used (Foll & Gaggiotti, 2008). The plotting and identification of outliers was performed with R software using the output of the MCMC algorithm.

A genotype accumulation curve was drawn with the package poppr in Rstudio v3.5.2 to determine the minimum number of loci necessary to discriminate between individuals in a population. This function randomly sampled the loci without replacement and counted the number of multilocus genotypes observed.

Estimates of number of alleles (N_a_), number of private alleles (N_ap_), mean observed heterozygosity (H_o_), mean genetic diversity (H_s_) and estimate of F_is_ following Nei (1987) were obtained using the package hierfstat of Rstudio v.5.2.

The Polymorphism Information Content (PIC) of each microsatellite locus was evaluated using the formula of Botstein & al. (1980), with n corresponding to the number of alleles, p_i_ to the frequency of the i^th^ allele, and p_j_ to the frequency of the j^th^ allele:

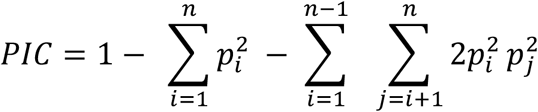

Markers with PIC value greater than 0.5 were considered highly informative (Botstein & al., 1980).

#### *Population structure, molecular variance and diversity analyses

Population structure was analysed on MLGs with the Bayesian model-based clustering software STRUCTURE v.2.3.4 (Pritchard & al., 2000). Simulations were performed under an admixture model with independent allele frequencies. The analysis was carried out for a number of tested clusters (K) ranging from 1 to 20, 10 runs, 100,000 burn-in period and 100,000 Markov Chain Monte Carlo (MCMC) repetitions after burning. The most likely number of populations (K) was identified using the Delta K method implemented in Structure Harvester. Structure results were summarized using Excel to obtain the probability of each lineage to belong to each cluster. Lineages of the 14 populations were assigned to one genetic cluster, or several genetic clusters when genotypes were admixed. A lineage was assigned to a genetic group if > 50% of its genome fraction value was derived from that group.

A Minimum Spanning Network on MLGs based on Nei distances with 1000 bootstraps and neighbor joining clustering method was performed using the package poppr (kamvar & al., 2014) of R Studio V3.5.2.

To assess the partitioning of the total genetic variance within and among populations, an analysis of molecular variance (AMOVA) based on *F*-statistics with 999 permutations was performed using GenAlEx 6.5 (Peakall & Smouse, 2012).

The genetic distance between populations was assessed by calculating pairwise F_st_ using the software Arlequin v3.5.2.2 (Excoffier & Lischer, 2010).

Diversity measures incorporate both genotypic richness R (number of genotypes in a population; Dorken & Eckert, 2001) and the Eveness index E_5_ (distribution of genotypes within a population; Grünwald & al., 2003). Common diversity indices were calculated, i.e. Shannon Weaver (H), Stoddart and Taylor’s index (G) and Simpson’s index (lambda). Finally, several genetic diversity parameters, including the number of alleles per population (Na), the observed heterozygosity (H_o_), the expected heterozygosity (H_e_), the number of multilocus genotypes (MLGs), the number of expected MLG at the smallest sample size ≥ 10 based on rarefaction (eMLGs), and Weir and Cockerham estimates of F statistics over all loci (F_st_), were estimated using the software Genclone v 2.0 (Arnaud-Haond & Belkhir, 2007), the packages poppr (Kamvar & al., 2014) and Hierfstat (Goudet, 2005) of Rstudio v3.5.2.

#### *Phylogenetic analysis

Several international gene banks have accessions from durum wheat landraces collected through the 20^th^ century in their collections. We ordered from the U.S. National Plant Germplasm System (NPGS) of the USDA (https://npgsweb.ars-grin.gov/) seed samples corresponding to 39 accessions from durum wheat landraces carrying the names ‘Bidi’, ‘Chili’ and ‘Mahmoudi’ from Tunisia, Algeria, Morocco and Italy (Table-ESM2). A 40^th^ accession, called ‘Mahmoudi-101’, was provided by the NGBT. Four individuals of each of the 40 accessions were genotyped with the nine SSR markers as described previously. In addition, three different seed lots were collected from the modern variety ‘Karim’ provided by the NGBT, the CRP-Wheat Septoria Phenotyping Platform, and the National Agronomic Institute of Tunisia (INAT). 30, 30 and 12 individuals from each respective lot were genotyped. Then, a phylogenetic analysis was conducted integrating genotypes of the 14 studied populations, the 40 accessions from landraces and the three seed lots from ‘Karim’. A phylogenetic tree based on Neighbour Joining clustering method on Edward’s distances, with 1000 bootstraps, was generated using the package poppr of Rstudio V3.5.2.

### Evaluation of phenotypic characters

#### *Determination of the ploidy level in lineages

Preliminary molecular and phenotypic analyses revealed lineages within eight populations belonging to the same genetic groups and being phenotypically more similar to hexaploid wheat species rather than tetraploid durum wheat. The ploidy level of 14 lineages, of the landrace Mahmoudi-101 and of variety Karim was verified by establishing their karyotype. The seeds were sterilized 10 min with sodium hypochlorite 5% and then rinsed with water. They were placed for 48 hours at 4°C in the dark for germination on a petri dish containing a water-moistened filter paper and then moved at 22°C in an oven for 48 hours. Roots of 2 cm were cut and placed in a glass tube containing a solution of ultra-pure water previously cooled for 24 hours on ice to stop mitosis. The roots were then fixed in an ethanol/acetic acid solution (3:1) at 4°C for 24 hours. The roots were placed in 1 mL of acetocarmine (carmin 10 g/L with 45% acetic acid) in a watch glass for 1 hour. The tip of a meristem is placed on a slide with a drop of 45% acetic acid, covered with a coverslip and then taped gently with the tip of a pencil to release the cells. The slide is lightly heated for a few seconds at 90 °C to remove the cytoplasmic veil. The quality of the spreads chromosomes is observed under a conventional phase contrast microscope. The photos are taken in bright-field at X40 magnification. Chromosome numbers are counted for five cells to estimate the hexaploid/tetraploid nature of the plants.

All hexaploid lineages were removed from further analyses on genetic and phenotypic diversity in the 14 populations.

#### *Agro-morphological characterization of landraces

Four representative spikes were randomly sampled at maturity from each lineage sown as a headrow in the Kodia Bou Salem’s field trial. Fifteen qualitative and quantitative agro-morphological traits were measured on the plants in the field or on the sampled spikes: plant height (HI), heading date (HS), spike shape (SS), spike colour (SC), spike density (SD), length of awns in relation to spike (LAS), length of spike without awns (LSWA), length of awns (LA), awn colour (AC), anthocyanin pigmentation of awns (PgA), number of spikelets per spike (NSS), colour of grains (CG), shape of grains (SG), and thousand grain weight (TGW). Scoring was done according to the recommendations of the International Plant Genetic Resource Institute (IPGRI) and the International Union for the Protection of New Varieties of Plants (UPOV) wheat descriptor lists (Table ESM3). Some lineages were lost during the field trial leaving us with 273 lineages for which a complete genotypic and phenotypic data set was available.

#### *Evaluation of landraces for resistance to *Septoria tritici* blotch

During the trial conducted on the CRP-Wheat Septoria Phenotyping Platform at Kodia Bou Salem, lineages from landraces and reference varieties were evaluated for their resistance to *Septoria tritici* blotch (STB). In the field, spreader rows of the modern variety ‘Karim’, highly deployed in Tunisia and considered susceptible (Berraies & al., 2014), were sown in all blocks perpendicularly to head-rows. Spreader rows and head-rows were spray-inoculated with the *Zymoseptoria tritici* strain IPO91009, collected in Tunisia (Béja) in 1991 (Kema & al., 1996). The inoculation was performed twice on 15 and 29 January 2018, between Zadoks’ 13 and (three fully unfolded leaves) and Zadoks’ 26 (tillering) stages with a conidial suspension adjusted to the concentration 10^6^ spores.mL^−1^. STB severity was assessed at two different dates following the “double digit” scoring method (Saari & Prescott, 1975). The area under the disease progression curve (AUDPC) was calculated from the percentage of disease severity (DS) at both observation dates (Sharma & Duveiller, 2007; Das & al., 1992) according to the formula:

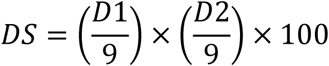

where, D1 is the first digit (vertical disease progress) and D2 is the second digit (severity of infection), and

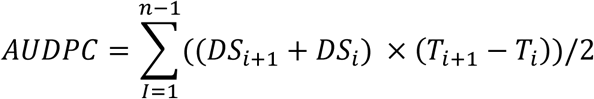

where, DS_i_ is disease severity at the i^th^ assessment, T_i_ is the time (number of days) on which the i^th^ assessment was performed, and n is the total number of assessments.

Statistical analyses were performed to assess differences between populations for their resistance/susceptibility to STB at p-value=0.05. Kruskall-Wallis tests were applied and Mann–Whitney tests were performed using Rstudio v 3.5.2.

#### *Statistical analysis of quantitative traits

A matrix of correlations between the nine quantitative traits was calculated at confidence interval of 0.99 using the package corrplot of Rstudio v 3.5.2. One-way Anova and Kruskall-Wallis tests – parametric and non-parametric tests, respectively – were applied depending on the trait’s distribution to test for phenotypic differences between populations. For the non-normally distributed traits, a first normalization was performed using the boxcox function (Box & Cox, 1964) from the package MASS (Venables & Ripley, 2002). In case a significant difference was detected for a trait, post-hoc tests were performed using Rstudio v 3.5.2: a Tukey’s Honest Significant Difference test (*i*.*e*. parametric) or a Mann–Whitney test (*i*.*e*. non-parametric), at p-value=0.001.

#### *Phenotypic diversity between populations and genetic groups

For qualitative phenotypic traits, we considered the most frequent phenotypic class among the four spikes of each lineage. For quantitative traits, the mean values of the four spikes were categorized into classes (Table ESM3). All qualitative and quantitative values grouped into classes were used to calculate the Shannon–Weaver diversity index (H) (Shannon & Weaver 1949; Jain & al. 1975) according to the formula:

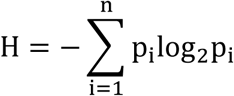

where p_i_ is the frequency of individuals from the i^th^ class and n is the number of classes for the designated phenotypic trait.

A relative phenotypic diversity index (H’) was calculated according to the formula:

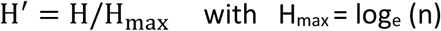

#### *Factor Analysis of Mixed Data (FAMD)

A Factor Analysis of Mixed Data (FAMD) was performed using the FactoMineR and factoextra packages in Rstudio v3.5.2, using both quantitative and qualitative variables to ensure a balance of their influence in the analysis. This method allows to reveal similarities between lineages and to explore associations between all variables.

### Relationship between genotypic and phenotypic diversity parameters

A P_st_-F_st_ analysis was performed using the 273 lineages for which both phenotypic and genetic data were available to test whether phenotypic differences in nine quantitative traits between populations were due to selection or genetic drift. P_st_ was used as an approximation of Q_st_, whose exact calculation requires common garden experiments to measure the additive genetic variances (Leinonen & al., 2008; Pujol & al., 2008; Brommer, 2011) and was not possible here.

The genetic differentiation F_st_ and the corresponding 95% confidence interval (CI) were calculated using the package hierfstat. P_st_ was estimated according to Brommer (2011) according to the formula:

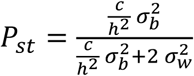

where 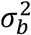 and 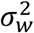 are the phenotypic variances between and within populations, respectively, c is an estimate of the proportion of the total variance due to additive genetic effects across populations, and h^2^ is the heritability (the proportion of phenotypic variance due to additive genetic effects). When P_st_=F_st_, the differentiation of quantitative traits may be the result of genetic drift, even if a contribution of natural selection cannot be discarded nor estimated. If P_st_>F_st_, quantitative traits have a higher level of differentiation, which could be an evidence for directional selection (or heterogeneous selection). Finally, if P_st_<F_st_, quantitative traits are less diversified than neutral differentiation, suggesting that these traits have been under the influence of stabilizing selection (or homogeneous selection).

As the determination of c is difficult, a sensitive analysis was performed using the package Pstat of R studio v3.5.2 to infer the robustness of the estimation of c/h^2^ which is critical for P_st_ to correctly approximate Q_st_. The lower critical c/h^2^ ratio is obtained when P_st_ exceeds F_st_ and at null assumption (c/h^2^ = 1) the proportion of phenotypic variance due to additive genetic effects is the same within and across populations. The trait is considered divergent at the point where P_st_ exceeds F_st_ (c<h^2^). P_st_ values were extracted at null assumption and P_st_ with a 95% CI were estimated using Pstat package with 1000 permutations. If the lower 95% CI of P_st_ is higher than F_st_ then the quantitative trait is considered highly divergent. For each trait with P_st_ > F_st_, the robustness of the P_st_ as an approximation of F_st_, indicating local adaptation, was estimated by the critical c/h^2^ value. This ratio is referring to the value when the lower 95% CI of P_st_ equal the upper 95% CI of F_st_ as described in Brommer & al. (2011). As the critical c/h^2^ was here lower than 0.20 for some traits, comparing P_st_ and F_st_ allowed us to make robust conclusions on the selection regime (Brommer, 2011). The critical c/h^2^ ratio was calculated according to the formula:

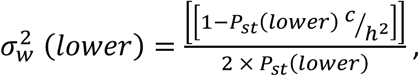

Then:

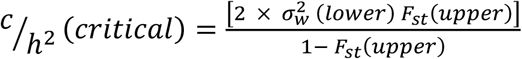

## Acknowledgments

This study was supported by grants from INRAE department Plant Health and Environment, for the APÔGÉ Project and for the PhD thesis of Safa Ben Krima, covering the 2017-2020 period. We thank Sonia Hamza (INAT, Tunisia) for assistance at the beginning of the project and for providing seeds of the durum wheat cultivar Karim, Marie-Hélène Muller (INRAE AGAP) for her assistance with population genetics analyses, and Anne-Lise Boixel and Florence Carpentier (INRAE BIOGER) for assistance with statistical analyses. We are grateful with the genebanks from USDA ARS and from CIMMYT for providing historical seed samples from Tunisian durum wheat landraces. And finally, we would like to express our deep gratitude to Tunisian farmers who always welcomed us and allowed us to work with their familial heritage, their durum wheat landraces.

## References

Agnoun Y, Yelome I, Sié M, Albar L, Ghesquière A, Silue D. 2019. Resistance of selected Oryza glaberrima landraces and their intra-specific breeding lines to Beninese rice yellow mottle virus isolates. Crop Protection 119:172–176. DOI: 10.1016/j.cropro.2019.01.022

Akem C, Ceccarelli S, Erskine W, Lenné J. 2000. Using genetic diversity for disease resistance in agricultural production. Outlook on Agriculture 29:25–25. DOI: 10.5367/000000000101293013

Al Khanjari S, Filatenko A, Hammer K, Buerkert A. 2008. Morphological spike diversity of Omani wheat. Genetic Resources and Crop Evolution 55: 1185–1195. DOI: 10.1007/s10722-008-9319-9

Alemu A, Feyissa T, Letta T, Abeyo B. 2020. Genetic diversity and population structure analysis based on the high-density SNP markers in Ethiopian durum wheat (Triticum turgidum ssp. durum). BMC Genetics 21: 18. DOI: 10.1186/s12863-020-0825-x

Ammar K, Gharbi MS, Deghaies M. 2011. Wheat in Tunisia. In: Bonjean A, Angus WM & Van Ginkel M (Eds). The world wheat book, A history of wheat breeding, Volume 2. Lavoisier Publishing, Paris.

Arnaud-Haond S, Belkhir K. 2007. GENCLONE: a computer program to analyse genotypic data, test for clonality and describe spatial clonal organization. Molecular Ecology Notes 7: 15–17. DOI: 10.1111/j.1471-8286.2006.01522.x

Asmamaw M, Keneni G, Kassahun T. 2019. Genetic diversity of Ethiopian durum wheat (Triticum durum Desf.) landrace collections as revealed by SSR Markers. Advances in Crop Science and Technology 7(1): 1000413. DOI: 10.4172/2329-8863.1000413

Ayadi S, Karmous C, Hammami Z, Tamani N, Trifa Y, Esposito S, Rezgui S. 2012. Genetic variability of nitrogen use efficiency components in Tunisian improved genotypes and landraces of durum wheat. Agricultural Science Research Journals 2(11): 591–601. DOI: 10.13140/RG.2.1.4907.2401

Ayed S, Karmous C, Trifa Y, Slama-Ayed O, Slim-Amara H. 2010. Phenotypic diversity of Tunisian durum wheat landraces. African Crop Science Journal 18(1): 35–42. DOI: 10.4314/acsj.v18i1.54197

Babay E, Mnasri SR, Mzid R, Ben Naceur M, Hanana M. 2019. Quality selection and genetic diversity of Tunisian durum wheat varieties using SSR markers. Bioscience Journal 35(4): 1002–1012. DOI: 10.14393/BJ-v35n4a2019-42301

Bechere E, Belay G, Mitiku D, Merker A. 1996. Phenotypic diversity of tetraploid wheat landraces from northern and north-central regions of Ethiopia. Hereditas 124: 124–172. DOI: 10.1111/j.1601-5223.1996.00165.x

Beharav A, Golan G, Levy A. 1997. Evaluation and variation in response to infection with Puccinia striiformis and Puccinia recondita of local wheat landraces. Euphytica 94: 287–293. DOI: 10.1023/A:1002983824125

Belaid A. Durum wheat in WANA: Production, trade, and gains from technological change. 2000. In: Royo C, Nachit M, Di Fonzo N, Araus JL (Eds). Seminar on durum wheat improvement in the Mediterranean region: New challenges. Zaragoza (Spain), CIHEAM, pp 35–49. URL: http://om.ciheam.org/om/pdf/a40/00600004.pdf

Belhadj H, Medini M, Bouhaouel I, Amara H. 2015. Analyse de la diversité phénotypique de quelques accessions autochtones de blé dur (Triticum turgidum ssp. durum Desf.) du sud tunisien. Journal of new science 24(5): 1115–1125. URL: https://www.jnsciences.org/agri-biotech/32-volume-24.html

Bellon MR. 1996. The dynamics of crop infraspecific diversity: A conceptual framework at the farmer level. Economic Botany 50: 26–39. DOI: 10.1007/BF02862110

Berraies S, Ammar K, Gharbi M, Yahyaoui A, Rezgui S. 2014. Quantitative inheritance of resistance to Septoria tritici blotch in durum wheat in Tunisia. Chilean journal of agricultural research 74: 35–40. DOI: 10.4067/S0718-58392014000100006

Bœuf F. 1926. Amélioration de la culture du Blé en Tunisie. Revue de botanique appliquée et d’agriculture coloniale 63 (6^è^me année): 657–666. DOI: 10.3406/jatba.1926.4456

Botstein D, White RL, Skolnic M, Davis RW. 1980. Construction of a genetic linkage map in man using restriction fragment length polymorphisms. American journal of human genetics 32(3): 314–331. PMID: 6247908

Bouacha OD, Rezgui S. 2019. Spaghetti quality: Comparison between landraces and high yielding Tunisian durum wheat varieties. Journal of New Sciences 64(7): 4056–4060. E-ISSN: 2286-5314

Box GEP, Cox DR. 1964. An analysis of transformations. Journal of the Royal Statistical Society. Series B (Methodological) 26(2): 211–252. URL: https://www.jstor.org/stable/2984418

Brommer JE. 2011. Whither Pst? The approximation of Qst by Pst in evolutionary and conservation biology. Journal of Evolutionary Biology 24(6): 1160–1168. DOI: 10.1111/j.1420-9101.2011.02268.x

Brush StB, Meng E. 1998. Farmers’ valuation and conservation of crop genetic resources. Genetic Resources and Crop Evolution 45: 139–150. DOI: 10.1023/A:1008650819946

Chamekh Z, Karmous C, Ayadi S, Sahli A, Hammami Z, Fraj MB, Benaissa N, Trifa Y, Slim-Amara H. 2015. Stability analysis of yield component traits in 25 durum wheat (Triticum durum Desf.) genotypes under contrasting irrigation water salinity. Agricultural Water Management 152: 1–6. DOI: 10.1016/j.agwat.2014.12.009

Conversa G, Lazzizera C, Bonasia A, Cifarelli S, Losavio F, Sonnante G, Elia A. 2020. Exploring on-farm agro-biodiversity: a study case of vegetable landraces from Puglia region (Italy). Biodiversity and Conservation 29: 747–770. DOI: 10.1007/s10531-019-01908-3

Das MK, Rajaram S, Mundt CC, Kronstad WE. 1992. Inheritance of slow-rusting resistance to leaf rust in wheat. Crop science 32(6): 1452–1456. DOI: 10.2135/cropsci1992.0011183X003200060028x

Deghais M, Kouki M, Gharbi MS, El Felah M. 2003. Les variétés de céréales cultivées en Tunisie: blé dur, blé tendre, orge et triticale. INRAT. 447p.

Deghais M, Kouki M, Gharbi MS, El Felah M. 2007. Les variétés de céréales cultivées en Tunisie: blé dur, blé tendre, orge et triticale. INRAT. 445p.

De Luca D, Cennamo P, Del Guacchio E, Di Novella R, Caputo P. 2018. Conservation and genetic characterisation of common bean landraces from Cilento region (southern Italy): high differentiation in spite of low genetic diversity. Genetica 146: 29–44. DOI: 10.1007/s10709-017-9994-6, PMID: 29030763

De Ron AM, Bebeli PJ, Negri V, Vaz Patto MC, Revilla P. 2018. Warm season grain legume landraces from the South of Europe for germplasm conservation and genetic improvement. Frontiers in Plant Science 9: p1524. DOI: 10.3389/fpls.2018.01524, PMID: 30405662

Dorken ME, Eckert CG. 2001. Severely reduced sexual reproduction in northern populations of a clonal plant, Decodon verticillatus (Lythraceae). Journal of Ecology 89: 339–350. DOI: 10.1046/j.1365-2745.2001.00558.x

Ehdaie B, Waines JG, Hall AE. 1988. Differential responses of landrace and improved spring wheat genotypes to stress environments. Crop Science 28: 838–842. DOI: 10.2135/cropsci1988.0011183X002800050024x

Erroux J. 1991. « Blé ». In : Encyclopédie berbère, 10, 1991, document B81, mise en ligne le 01 mai 2013, consultée le 01 mars 2020. URL: http://journals.openedition.org/encyclopedieberbere/1766

Excoffier L, Lischer HEL. 2010. Arlequin suite ver 3.5: A new series of programs to perform population genetics analyses under Linux and Windows. Molecular Ecology Resources 10(3): 564–567. DOI: 10.1111/j.1755-0998.2010.02847.x, PMID: 21565059

Feldman M. 2001. Origin of cultivated wheat. In: Bonjean AP, Angus WJ (Eds). The world wheat book, A history of wheat breeding. Lavoisier Publishing, Paris, pp 3–56.

Ferjaoui S, Sbei A, Aouadi N, Hamza S. 2011. Monogenic inheritance of resistance to Septoria tritici blotch in durum wheat ‘Agili’. International Journal of Plant Breeding 5: 17–20.

Ferjaoui S, M’Barek SB, Bahri B, Slimane RB, Hamza S. 2015. Identification of resistance sources to Septoria tritici blotch in old tunisian durum wheat germplasm applied for the analysis of the Zymoseptoria tritici-durum wheat interaction. Journal of Plant Pathology 97(3): 471–481. DOI: 10.4454/JPP.V97I3.028

Figliuolo G, Mazzeo M, Greco I. 2007. Temporal variation of diversity in Italian durum wheat germplasm. Genetic Resources and Crop Evolution 54(3): 615–626. DOI: 10.1007/s10722-006-0019-z

Foll M, Gaggiotti O. 2008. A genome scan method to identify selected loci appropriate for both dominant and codominant markers: a Bayesian perspective. Genetics 180: 977–993. DOI: 10.1534/genetics.108.092221, PMID: 18780740

Gautier A, Marcel TC, Confais J, Crane C, Kema G, Suffert F, Walker A-S. 2014. Development of a rapid multiplex SSR genotyping method to study populations of the fungal plant pathogen Zymoseptoria tritici. BMC Research Notes 7: 373. DOI: 10.1186/1756-0500-7-373

Gharbi MS, Elfelah M. 2013. Les céréales en Tunisie: plus d’un siècle de recherche variétale. Annales de l’INRAT 86 (Numéro Spécial Centenaire): 45–68. ISSN: 0365-4761

Goudet J. 2005. Hierfstat, a package for R to compute and test variance components and F-statistics. Molecular Ecology Notes 5: 184–186. DOI: 10.1111/j.1471-8278.2004.00828.x

Govindaraj M, Vetriventhan M, Srinivasan M. 2015. Importance of genetic diversity assessment in crop plants and its recent advances: an overview of its analytical perspectives. Genetics Research International 2015: Article ID 431487. DOI: 10.1155/2015/431487, PMID: 25874132

Grünwald NJ, Goodwin SB, Milgroom MG, Fry WE. 2003. Analysis of genotypic diversity data for populations of microorganisms. Phytopathology 93(6): 738–746. DOI: 10.1094/PHYTO.2003.93.6.738, PMID: 18943061

Hammami R, Sissons M. 2020. Durum wheat products, couscous. In: Igrejas G, Ikeda T, Guzmán C (Eds). Wheat quality for improving processing and human health. Springer, Cham. DOI: 10.1007/978-3-030-34163-3_15

Hammer K, Diederichsen A. 2009. Evolution, status and perspectives for landraces in Europe. In: Vetelainen M, Negri V, Maxted N (Eds). European landraces: on-farm conservation, management and use. Bioversity Technical Bulletin No. 15, Bioversity International publisher, Rome, Italy, pp 23–43. URL: https://www.bioversityinternational.org/e-library/publications/detail/european-landraces-on-farm-conservation-management-and-use/

Huhn MR, Elias EM, Ghavami F, Kianian SF, Chao S, Zhong S, Alamri MS, Yahyaoui A, Mergoum M. 2012. Tetraploid Tunisian wheat germplasm as a new source of Fusarium head blight resistance. Crop Science 52(1): 136–145. DOI: 10.2135/cropsci2011.05.0263

Hurtado P, Olsen K, Buitrago C, Ospina C, Marin J, Duque M, Wongtiem P, Wenzel P, Killian A, Adeleke M, Fregene M. 2008. Comparison of simple sequence repeat (SSR) and diversity array technology (DArT) markers for assessing genetic diversity in cassava (Manihot esculenta Crantz). Plant Genetic Resources 6(3): 208–214. DOI: 10.1017/S1479262108994181

Jain SK, Qualset CO, Bhatt GM, Wu KK. 1975. Geographical patterns of phenotypic diversity in a word collection of durum wheats. Crop Science 15: 700–704. DOI: 10.2135/cropsci1975.0011183X001500050026x

Kabbaj H, Sall AT, Al-Abdallat A, Geleta M, Amri A, Filali-Maltouf A, Belkadi B, Ortiz R, Bassi FM. (2017). Genetic diversity within a global panel of durum wheat (Triticum durum) landraces and modern germplasm reveals the history of alleles exchange. Frontiers in Plant Science 8: 1277. DOI: 10.3389/fpls.2017.01277, PMID: 28769970

Kamvar ZN, Tabima JF, Grünwald NJ. 2014. Poppr: an R package for genetic analysis of populations with clonal, partially clonal, and/or sexual reproduction. PeerJ 2: e281. DOI: 10.7717/peerj.281, PMID: 24688859

Kema GHJ, Annone JG, Sayoud R, van Silfhout CH, van Ginkel M, de Bree J. 1996. Genetic variation for virulence and resistance in the wheat-Mycosphaerella graminicola pathosystem. I. Interactions between pathogen isolates and host cultivars. Phytopathology 86: 200–212. DOI: 10.1094/Phyto-86-200

Knezevic D, Paunovic A, Madic M, Djukic N. 2007. Genetic analysis of nitrogen accumulation in four wheat cultivars and their hybrids. Cereal Research Communications 35(2): 633–636. Proceedings of the VI. Alps-Adria Scientific Workshop, Obervellach, Austria, 30 April–5 May 2007. DOI: 10.1556/CRC.35.2007.2.117

Korzun V, Börner A, Worland AJ, Law CN, Röder MS. 1997. Application of microsatellite markers to distinguish inter-varietal chromosome substitution lines of wheat (Triticum aestivum L.). Euphytica 95: 149–155. DOI: 10.1023/A:1002922706905

Kuckuck H, Kobabe G, Wenzel G. 1991. Fundamentals of Plant Breeding. 1st Edition. Springer-Verlag Berlin Heidelberg, Germany, IX 236p. ISBN: 978-3-642-75394-7

Kyratzis A, Nikoloudakis N, Katsiotis A. 2019. Genetic variability in landraces populations and the risk to lose genetic variation. The example of landrace ‘Kyperounda’ and its implications for ex situ conservation. PLoS ONE 14(10): e0224255. DOI: 10.1371/journal.pone.0224255

La Rovere R, Thabet C, Ammar K, Sferi R. 2010. The Tunisian wheat sector in the new liberalization scenario. New Medit 9(1): 13–23.

Laumont P, Erroux J. 1962. Les blés tendres cultivés en Algérie. Annales de l’Institut national agronomique El Harrach 3: 1–60. URL: https://www.asjp.cerist.dz/en/article/15083

Leinonen T, O’Hara RB, Cano JM, Merilä J. 2008. Comparative studies of quantitative trait and neutral marker divergence: a meta-analysis. Journal of Evolutionary Biology 21: 1–17. DOI: 10.1111/j.1420-9101.2007.01445.x, PMID: 18028355

Leinonen T, McCairns R, O’Hara R, Merilä J. 2013. QST–FST comparisons: evolutionary and ecological insights from genomic heterogeneity. Nature Reviews Genetics 14: 179–190. DOI: 10.1038/nrg3395

Lovell DJ, Parker SR, Hunter T, Royle DJ, Coken RR. 1997. Influence of crop growth and structure on the risk of epidemics by Mycosphaerella graminicola (Septoria tritici) in winter wheat. Plant pathology 46: 126–138. DOI: 10.1046/j.1365-3059.1997.d01-206.x

MacKey J. 2005. Wheat, its concept, evolution andtaxonomy. In: Royo C, Nachit M, Di Fonzo N, Araus JL, Pfeiffer WH, Slafer GA (Eds). Durum wheat breeding: current approaches and future strategies, Vol. I. Howorth Press, New York, pp 3–62.

Martínez-Moreno F, Solís I, Noguero D, Blanco A, Özberk I, Nsarellah N, Elias E, Mylonas I, Soriano JM. (2020). Durum wheat in the Mediterranean Rim: historical evolution and genetic resources. Genetic Resources and Crop Evolution 67: 1415–1436. DOI: 10.1007/s10722-020-00913-8

Maxted N, Kell S, Toledo Á, Dulloo E, Heywood V, Hodgkin T, Hunter D, Guarino L, Jarvis A, Ford-Lloyd B. 2010. A global approach to crop wild relative conservation: Securing the gene pool for food and agriculture. Kew Bulletin 65: 561–576. DOI: 10.1007/s12225-011-9253-4

Medini M, Hamza S, Rebai A, Baum M. 2005. Analysis of genetic diversity in Tunisian durum wheat cultivars and related wild species by SSR and AFLP markers. Genetic Resources and Crop Evolution 52: 21–31. DOI: 10.1007/s10722-005-0225-0

Morris EK, Caruso T, Busco F, Fischer M, Hancock C, Maier TS, Meiners T, Müller C, Obermaier E, Prati D, Socher SA, Sonnemann I, Wäschke N, Wubet T, Wurst S, Rillig MC. 2014. Choosing and using diversity indices: insights for ecological applications from the German biodiversity exploratories. Ecology and evolution 4(18): 3514–3524. DOI: 10.1002/ece3.1155, PMID: 25478144

Nazco R, Villegas D, Ammar K, Peña RJ, Moragues M, Royo C. 2012. Can Mediterranean durum wheat landraces contribute to improved grain quality attributes in modern cultivars? Euphytica 185: 1–17. DOI: 10.1007/s10681-011-0588-6

Nazco R, Peña RJ, Ammar K, Villegas D, Crossa J, Moragues M, Royo C. 2014. Variability in glutenin subunit composition of Mediterranean durum wheat germplasm and its relationship with gluten strength. The Journal of Agricultural Science 152(3): 379–393. DOI: 10.1017/S0021859613000117, PMID: 24791017

Nefzaoui M, Udupa S, Gharbi M, Bouhadida M, Iraqi D.2014. Molecular diversity in Tunisian durum wheat accessions based on microsatellite markers analysis. Romanian Agricultural Research 31: 33–39. ISSN: 2067-5720

Negri V. 2003. Landraces in central Italy: where and why they are conserved and perspectives for their on-farm conservation. Genetic Resources and Crop Evolution 50: 871–885; DOI: 10.1023/A:1025933613279

Nei M. 1987. Molecular evolutionary genetics. Columbia University Press, New York.

Ouaja M, Aouini L, Bahri B, Ferjaoui S, Medini M, Marcel TC, Hamza S. 2020. Identification of valuable sources of resistance to Zymoseptoria tritici in the Tunisian durum wheat landraces. European Journal of Plant Pathology 156: 647–661. DOI: 10.1007/s10658-019-01914-9

Peakall R, Smouse PE. 2012. GenAlEx 6.5: genetic analysis in Excel. Population genetic software for teaching and research--an update. Bioinformatics (Oxford, England) 28(19): 2537–2539. DOI: 10.1093/bioinformatics/bts460

Perrings C. 2018. Conservation beyond protected areas: the challenge of landraces and crop wild relatives. In: Dayal V, Duraiappah A, Nawn N (Eds). Ecology, Economy and Society. Springer, Singapore. DOI: 10.1007/978-981-10-5675-8_8

Pielou EC. 1975. Ecological diversity. John Wiley and Sons. New York, pp 165.

Pietrusińska A, Monika Ż, Piechota U, Słowacki P, Moskal K. 2018. Searching for diseases resistance sources in old cultivars, landraces and wild relatives of cereals. A review. Annales UMCS, Agricultura LXXIII(4): 45–60. DOI: 10.24326/asx.2018.4.5

Portères R. 1958. Les appellations des céréales en Afrique (suite). Journal d’agriculture tropicale et de botanique appliquée 5(4-5): 311–364. DOI: 10.3406/jatba.1958.2469

Poudel D, Johnsen FH. 2009. Valuation of crop genetic resources in Kaski, Nepal: Farmers’ willingness to pay for rice landraces conservation. Journal of Environmental Management 90(1): 483–491. DOI: 10.1016/j.jenvman.2007.12.020, PMID: 18359142

Pritchard JK, Stephens M, Donnelly P. 2000. Inference of population structure using multilocus genotype data. Genetics 155: 945–959. PMCID: PMC1461096

Pujol B, Wilson AJ, Ross RIC, Pannell JR. 2008. Are Q (ST)-F-ST comparisons for natural populations meaningful? Molecular Ecology 17: 4782–4785. DOI: 10.1111/j.1365-294X.2008.03958.x, PMID: 19140971

Rawson HM. 1970. Spikelet number, its control and relation to yield per ear in wheat. Australian Journal of Biological Sciences 23: 1–16. DOI: 10.1071/BI9700001

Robbana C, Kehel Z, Ben Naceur M, Sansaloni C, Bassi F, Amri A. 2019. Genome-wide genetic diversity and population structure of Tunisian durum wheat landraces based on DArTseq technology. International Journal of Molecular Sciences 20(6): 1352. DOI: 10.3390/ijms20061352, PMID: 30889809

Röder MS, Korzun V, Wendehake K, Plaschke J, Tixier MH, Leroy P, Ganal MW. 1998. A microsatellite map of wheat. Genetics 149(4): 2007–2023. PMID: 9691054

Rosenberg NA, Mahajan S, Ramachandran S, Zhao C, Pritchard JK, Feldman MW. 2005. Clines, clusters, and the effect of study design on the inference of human population structure. PLoS Genetics 1(6): e70. DOI: 10.1371/journal.pgen.0010070

Royo C, Soriano JM, Alvaro F. 2017. Wheat: a crop in the bottom of the Mediterranean diet pyramid. In Mediterranean Identities – Environment, Society, Culture. InTech. DOI: 10.5772/intechopen.69184

Rufo R, Alvaro F, Royo C, Soriano JM. 2019. From landraces to improved cultivars: assessment of genetic diversity and population structure of Mediterranean wheat using SNP markers. PLoS ONE 14 (7): e0219867. DOI: 10.1371/journal.pone.0219867

Russell J, Fuller J, Macaulay M, Hatz BG, Jahoor A, Powell W, Waugh R. 1997. Direct comparison of levels of genetic variation among barley accessions detected by RFLPs, AFLPs, SSRs and RAPDs. Theoretical and Applied Genetics 95: 714–722. DOI: 10.1007/s001220050617

Saade ME. 1996. Adoption and impact of high yielding wheat varieties in Northern Tunisia. CIMMYT Economics Working Paper. 96–03. Mexico, D.F. ISSN: 0258-8587

Saari EE, Prescott JM. 1975. A scale for appraising the foliar intensity of wheat disease. Plant Disease Reporter 59: 377–380.

Sahri A, Chentoufi L, Arbaoui M, Ardisson M, Belqadi L, Birouk A, Roumet P, Muller MH. 2014. Towards a comprehensive characterization of durum wheat landraces in Moroccan traditional agrosystems: analysing genetic diversity in the light of geography, farmers’ taxonomy and tetraploid wheat domestication history. BMC evolutionary biology 14: 264. DOI: 10.1186/s12862-014-0264-2

Semagn K, Babu R, Hearne S, Olsen M. 2014. Single nucleotide polymorphism genotyping using Kompetitive Allele Specific PCR (KASP): overview of the technology and its application in crop improvement. Molecular Breeding 33: 1–14. DOI: 10.1007/s11032-013-9917-x

Shannon CE, Weaver W. 1949. The mathematical theory of communication. The University of Illinois. Urbana, Chicago, London. pp. 3–24.

Shannon CE. 2001. A mathematical theory of communication. ACM SIGMOBILE Mobile Computing and Communications Review 5(1): 3–55. DOI: 10.1145/584091.584093

Sharma RC, Duveiller E. 2007. Advancement toward new Septoria Blotch resistance wheats in south Asia. Crop Science 47: 961–968. DOI: 10.2135/cropsci2006.03.0201

Simpson EH. 1949. Measurement of diversity. Nature 163: 688. DOI: 10.1038/163688a0

Slim A, Ayed S, Slama-Ayed O, Robbana C, Jaime AT, Slim-Amara H. 2011. Morphological diversity of some qualitative traits in tetraploid wheat landrace populations collected in the South of Tunisia. International Journal of Plant Breeding 5(1): 67–70.

Slim A, Piarulli L, Chennaoui Kourda H, Rouaissi M, Robbana C, Chaabane R, Pignone D, Montemurro C, Mangini G. 2019. Genetic structure analysis of a collection of Tunisian durum wheat germplasm. International Journal of Molecular Sciences 20: 3362. DOI: 10.3390/ijms20133362, PMID: 31323925

Spitze K. 1993. Population structure in Dahpnia obtusa: quantitative genetic and allozymic variation. Genetics 135(2): 367–374. PMID: 8244001

Somers DJ, Isaac P, Edwards K. 2004. A high-density microsatellite consensus map for bread wheat (Triticum aestivum L.). Theoretical and Applied Genetics 109: 1105–1114. DOI: 10.1007/s00122-004-1740-7, PMID: 15490101

Soriano JM, Villegas D, Aranzana MJ, García del Moral LF, Royo C. 2016. Genetic structure of modern durum wheat cultivars and Mediterranean landraces matches with their agronomic performance. PLoS ONE 11: e0160983. DOI: 10.1371/journal.pone.0160983, PMID: 27513751

Sourdille P, Singh S, Cadalen T, Brown-Guedira GL, Gay G, Qi L, Gill BS, Dufour P, Murigneux A, Bernard M. 2004. Microsatellite-based deletion bin system for the establishment of genetic-physical map relationships in wheat (Triticum aestivum L.). Functional & Integrative Genomics 4: 12–25. DOI: 10.1007/s10142-004-0106-1, PMID: 15004738

Stoddart JA, Taylor JF. 1988. Genotypic diversity: Estimation and prediction in samples. Genetics 118: 705–711. PMID: 17246421

Targońska M, Bolibok-Brągoszewska H, Rakoczy-Trojanowska M.2016. Assessment of genetic diversity in secale cereale based on SSR markers. Plant Molecular Biology Reporter 34: 37–51. DOI: 10.1007/s11105-015-0896-4

Venables WN, Ripley BD. 2002. Modern Applied Statistics with S. Fourth Edition, Springer, New York. ISBN 0-387-95457-0, URL: http://www.stats.ox.ac.uk/pub/MASS4

Villa TCC, Maxted N, Scholten M, Ford-Lloyd B. 2007. Defining and identifying crop landraces. Plant Genetic Resources 3(3): 373–384. DOI: 10.1079/PGR200591

Wallace M, Bonhomme V, Russell J, Stillman E, George TS, Ramsay L, Wishart J, Timpany S, Bull H, Booth A, Martin P. 2019. Searching for the origins of bere barley: a geometric morphometric approach to cereal landrace recognition in archaeology. Journal of Archaeological Method and Theory 26: 1125–1142. DOI: 10.1007/s10816-018-9402-2

Weir BS, Cockerham CC. 1984. Estimating F-statistics for the analysis of population structure. Evolution 38(6): 1358–1370. DOI: 10.2307/2408641, PMID: 28563791

Xu XD, Feng J, Fan JR, Liu ZY, Li Q, Zhou YL, Ma ZH. 2018. Identification of the resistance gene to powdery mildew in Chinese wheat landrace Baiyouyantiao. Journal of Integrative Agriculture 17: 37–45. DOI: 10.1016/S2095-3119(16)61610-6

Yacoubi I, Nigro D, Sayar R, Masmoudi K, Seo YW, Brini F, Giove SL, Mangini G, Giancaspro A, Marcotuli I, Colasuonno P, Gadaleta A. 2020. New insight into the North-African durum wheat biodiversity: phenotypic variations for adaptive and agronomic traits. Genetic Resources and Crop Evolution 67: 445–455. DOI: 10.1007/s10722-019-00807-4

Yao F, Zhang X, Ye X, Li J, Long L, Yu C, Li J, Wang Y, Wu Y, Wang J, Jiang Q, Li W, Ma J, Wei Y, Zheng Y, Chen G. 2019. Characterization of molecular diversity and genome-wide association study of stripe rust resistance at the adult plant stage in Northern Chinese wheat landraces.BMC Genetics 20: Article number 38. DOI: 10.1186/s12863-019-0736-x, PMID: 30914040

Zeven AC, Waninge J. 1989. The presence of three groups of Scalavatis and other hexaploid bread wheat plants contaminating durum wheat fields in Cyprus.Euphytica 43: 117–124. DOI: 10.1007/BF00037904

Zeven AC. 1998. Landraces: a review of definitions and classifications. Euphytica 104: 127–139. DOI: 10.1023/A:1018683119237 Website: https://graingenes.org/GG3/

