## Supplementary Material for "Life story of Tunisian durum wheat landraces revealed by their genetic and phenotypic diversity"

### Supplementary Figures

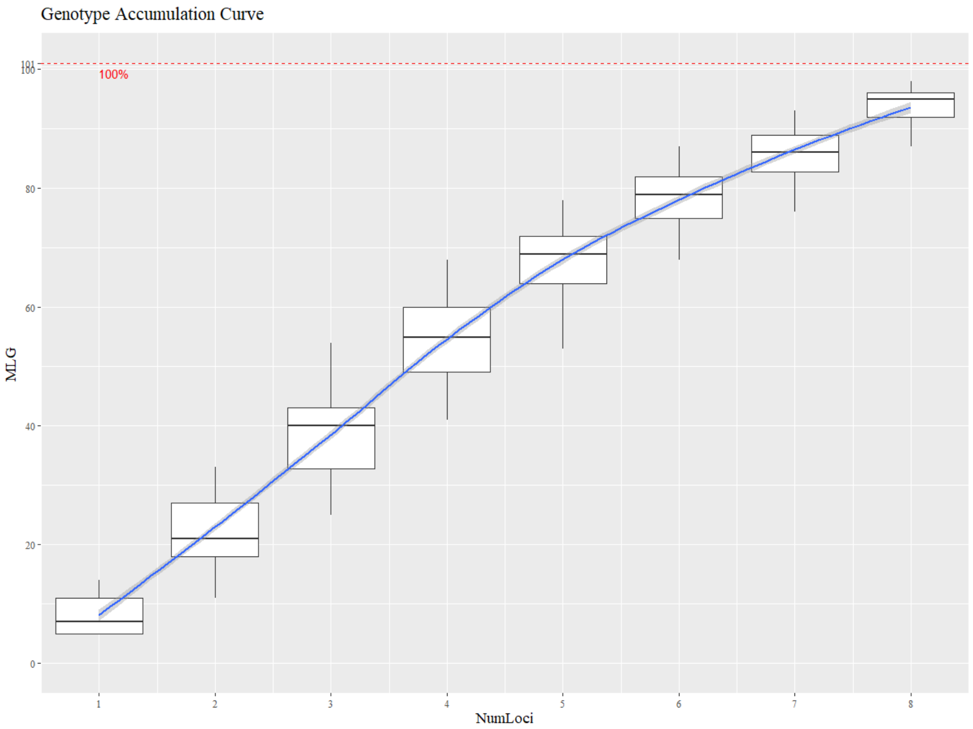


**Figure ESM1. Genotype accumulation curve for determining the minimum number of loci necessary to discriminate between individuals in a population.** This function randomly samples loci without replacement and count the number of multilocus genotypes observed.

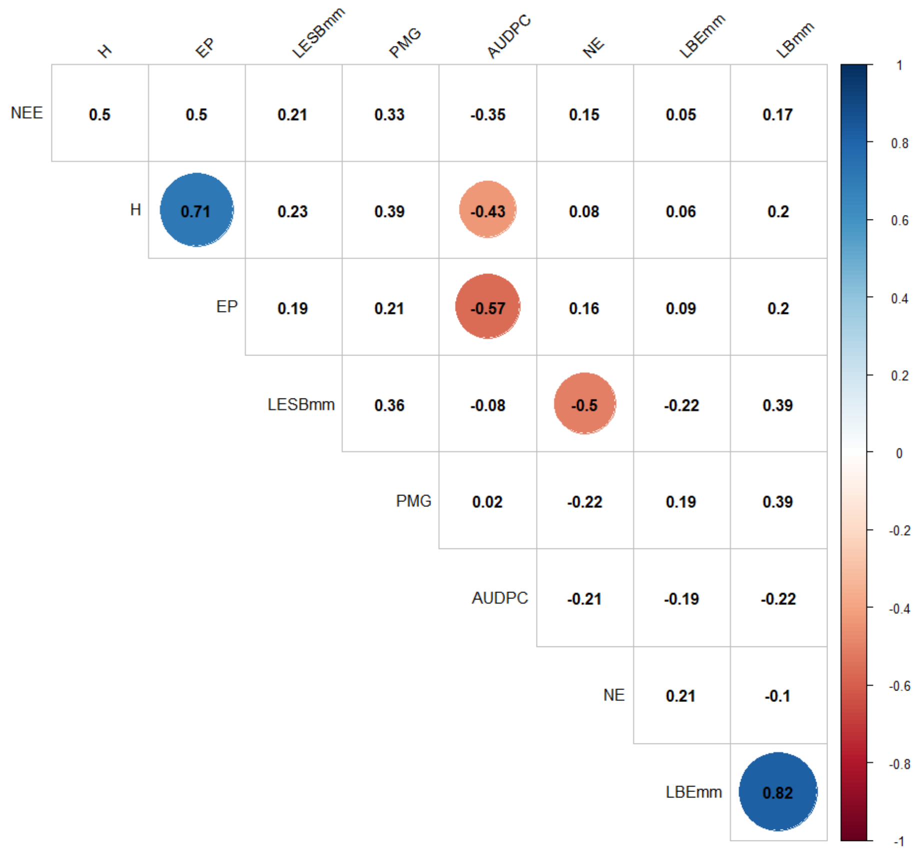


**Figure ESM2. Correlation plot between the 9 quantitative phenotypic traits (significant correlations at *p*< 0.01 are indicated by coloured circles).** HI: plant height; HS: heading date; AUDPC: area under disease progress curve; SD: spike density; LAS: length of awns in relation to spike; LSWA: length of spike without awns; LA: length of awns; NSS: number of spikelets per spike; TGW: thousand grains weight.

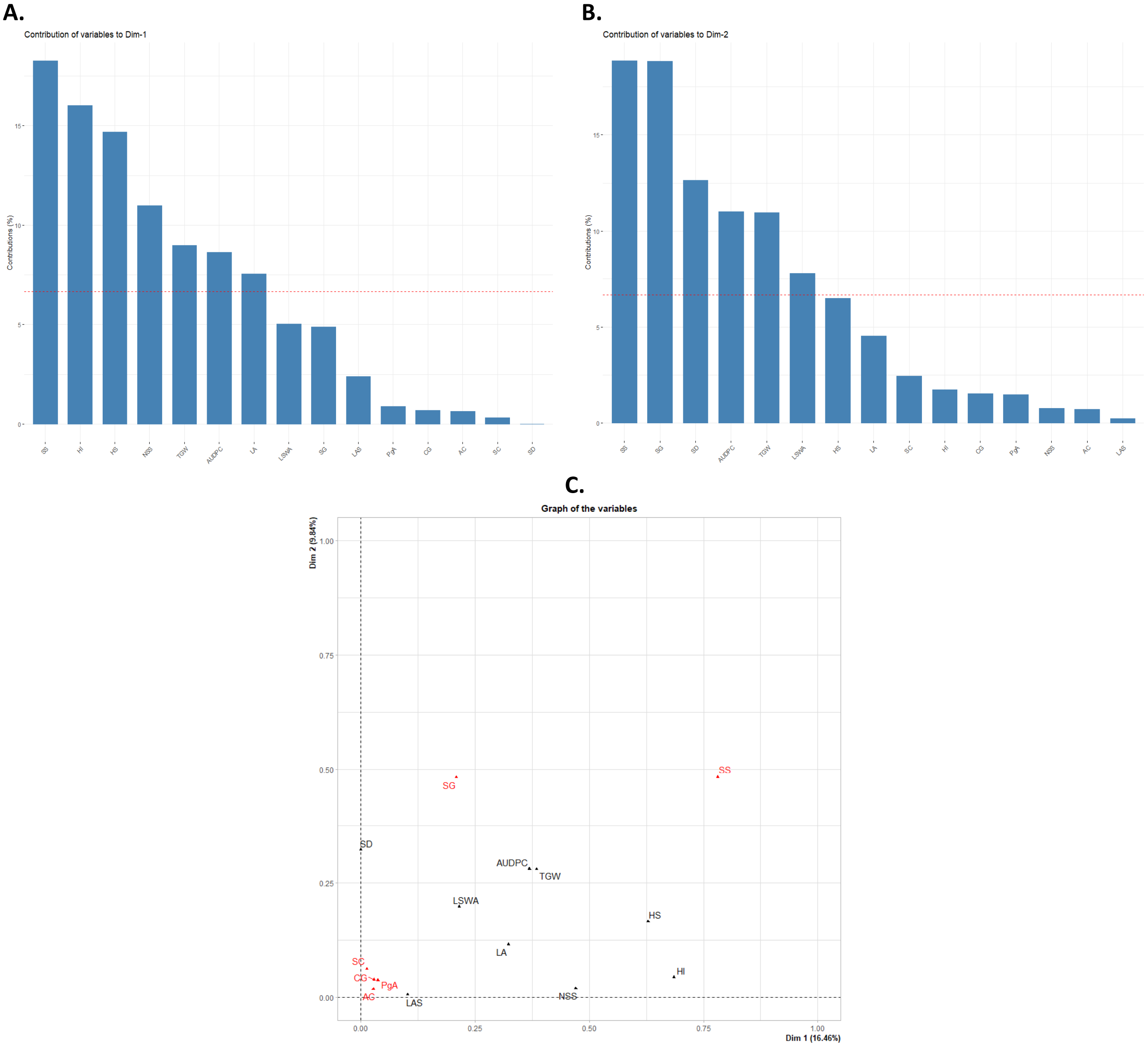


**Figure ESM3. FAMD output. A. Contribution of the different phenotypic traits to Dim1. B. Contribution of the different phenotypic traits to Dim2. C. Contribution of the different phenotypic traits to Dim1 and Dim2. Quantitative traits are labelled in black and qualitative traits are labelled in red.** HI: plant height; HS: heading date; AUDPC: area under disease progress curve; SD: spike density; LAS: length of awns in relation to spike; LSWA: length of spike without awns; LA: length of awns; NSS: number of spikelets per spike; TGW: thousand grains weight.

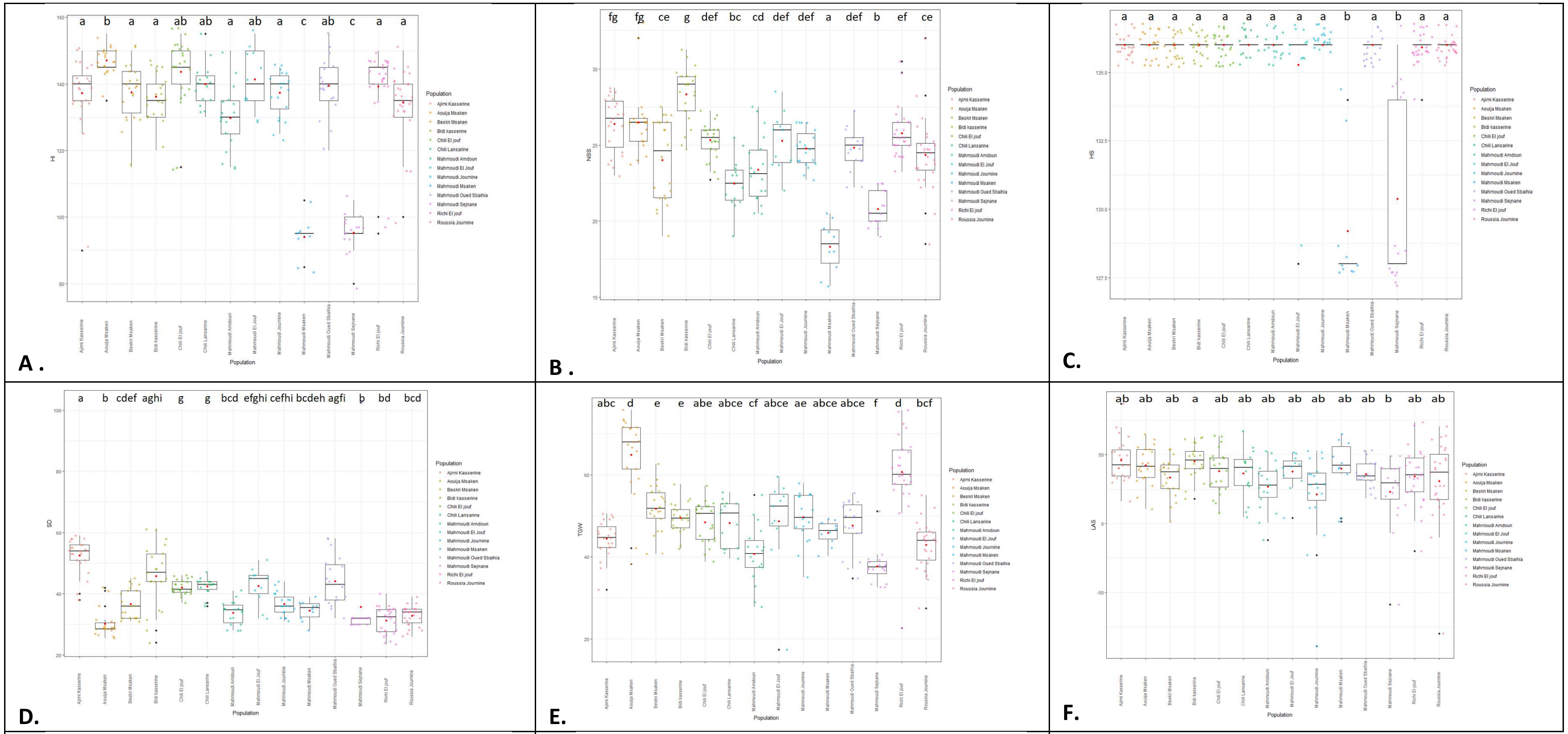


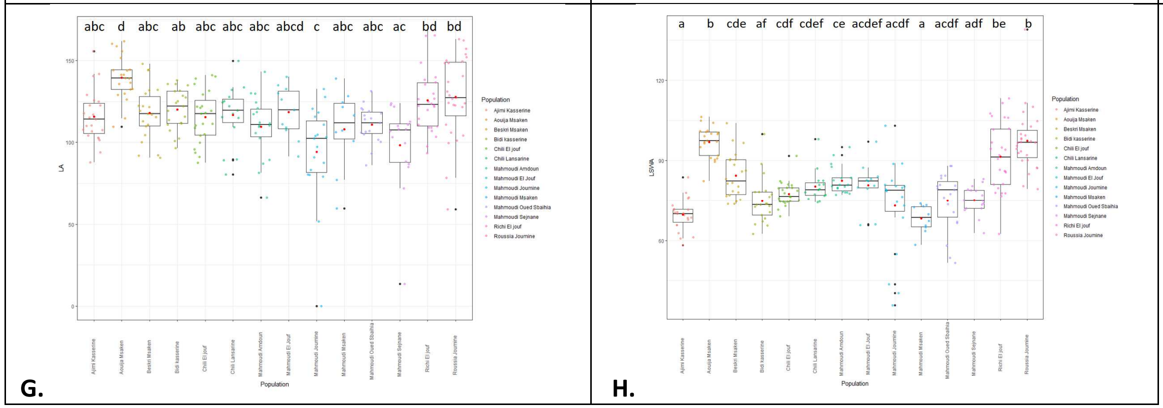


**Figure ESM4. Boxplots of quantitative phenotypic traits for the 14 durum wheat populations.** Means are represented by a red point. Populations with significantly different means are indicated by different letters after Kruskal-Wallis and Mann–Whitney tests at *p*=0.05. A. Plant height (HI); B. Number of spikelets per spike (NSS); C. Heading date (HS); D. Spike density (SD). E. Thousand grains weight (TGW); F. Length of awns in relation to spike (LAS); G. Length of awns (LA); H. Length of spike without awns (LSWA).

### Supplementary Tables

**Table ESM1. Matrix of $F_{st}$ between identified genetic groups (G1 to G7).**

| | **G1** | **G2** | **G3** | **G4** | **G5** | **G6** |
|---|---|---|---|---|---|---|
| **G2** | 0.404* | | | | | |
| **G3** | 0.485* | 0.573* | | | | |
| **G4** | 0.393* | 0.492* | 0.544* | | | |
| **G5** | 0.508* | 0.637* | 0.603* | 0.505* | | |
| **G6** | 0.524* | 0.628* | 0.562* | 0.553* | 0.645* | |
| **G7** | 0.357* | 0.455* | 0.457* | 0.471* | 0.556* | 0.602* |

*Significance of the $F_{st}$ at a threshold of 5%.

**Table-ESM2-Origin of Durum Wheat landraces stored at the USDA.**

| Names in the publication | Inventory USDA | Taxon | Collected from | Local name | Collectors |
|---|---|---|---|---|---|
| Mahmoudi (Cltr15501, 1972) | Cltr 15501 TR17ID SD | *Triticum turgidum* L. subsp. *durum* (Desf.) van Slageren | Tunisia, Tozeur | Mahmoudi | UN[1] |
| Bidi (Cltr3170, 1911) | Cltr 3170 TR13CA SD | *Triticum turgidum* L. subsp. *durum* (Desf.) van Slageren | Tunisia | Bidi AP 1 | Boeuf, F., BSV[2] |
| Bidi (Cltr3171, 1911) | Cltr 3171 TR15CA SD | *Triticum turgidum* L. subsp. *durum* (Desf.) van Slageren | Tunisia | Bidi AP 2 | Boeuf, F., BSV[2] |
| Bidi (Cltr3173, 1911) | Cltr 3173 TR13CA SD | *Triticum turgidum* L. subsp. *durum* (Desf.) van Slageren | Tunisia | Bidi AP 4 | Boeuf, F., BSV[2] |
| Bidi (Cltr3174, 1911) | Cltr 3174 TR13CA SD | *Triticum turgidum* L. subsp. *durum* (Desf.) van Slageren | Tunisia | Bidi AP 5 | Boeuf, F., BSV[2] |
| Mahmoudi (Cltr3233, 1911) | Cltr 3233 TR06ID SD | *Triticum turgidum* L. subsp. *durum* (Desf.) van Slageren | Tunisia | Mahmoudi Glabre AC 1 | Boeuf, F., BSV[2] |
| Mahmoudi (Cltr3234, 1911) | Cltr 3234 TR06ID SD | *Triticum turgidum* L. subsp. *durum* (Desf.) van Slageren | Tunisia | Mahmoudi Glabre AC 4 | Boeuf, F., BSV[2] |
| Mahmoudi (Cltr3235, 1911) | Cltr 3235 TR06ID SD | *Triticum turgidum* L. subsp. *durum* (Desf.) van Slageren | Tunisia | Mahmoudi Glabre AP 1 | Boeuf, F., BSV[2] |
| Mahmoudi (Cltr3236, 1911) | Cltr 3236 TR17ID SD | *Triticum turgidum* L. subsp. *durum* (Desf.) van Slageren | Tunisia | Mahmoudi Glabre AP 2 | Boeuf, F., BSV[2] |
| Mahmoudi (Cltr3237, 1911) | Cltr 3237 TR06ID SD | *Triticum turgidum* L. subsp. *durum* (Desf.) van Slageren | Tunisia | Mahmoudi Glabre AP 3 | Boeuf, F., BSV[2] |
| Mahmoudi (Cltr3241, 1911) | Cltr 3241 TR06ID SD | *Triticum turgidum* L. subsp. *durum* (Desf.) van Slageren | Tunisia | Mahmoudi Glabre RP 2 | Boeuf, F., BSV[2] |
| Mahmoudi (Cltr3242, 1911) | Cltr 3242 TR06ID SD | *Triticum turgidum* L. subsp. *durum* (Desf.) van Slageren | Tunisia | Mahmoudi Pubescent AP 1 | Boeuf, F., BSV[2] |
| Mahmoudi (Cltr3809, 1912) | Cltr 3809 TR13ID SD | *Triticum turgidum* L. subsp. *durum* (Desf.) van Slageren | Tunisia, Bizerte | Mahmoudi | ICMEW[3] in 1910 |
| Bidi (Cltr3811, 1912) | Cltr 3811 TR14CA SD | *Triticum turgidum* L. subsp. *durum* (Desf.) van Slageren | Tunisia, Siliana | Bidi | ICMEW[3] in 1910 |
| Mahmoudi (Cltr3816, 1912) | Cltr 3816 TR13CA SD | *Triticum turgidum* L. subsp. *durum* (Desf.) van Slageren | Tunisia, Nabeul | Mahmoudi | ICMEW[3] in 1910 |
| Mahmoudi (Cltr3824, 1912) | Cltr 3824 TR12ID SD | *Triticum turgidum* L. subsp. *durum* (Desf.) van Slageren | Tunisia, Beja | Mahmoudi | ICMEW[3] in 1910 |
| Mahmoudi (Cltr6876, 1923) | Cltr 6876 TR16ID SD | *Triticum turgidum* L. subsp. *durum* (Desf.) van Slageren | Tunisia | Mahmoudi | IGA[4] |
| Mahmoudi (PI150380, 1945) | PI 150380 TR06ID SD | *Triticum turgidum* L. subsp. *durum* (Desf.) van Slageren | Tunisia | Mahmoudi | Beall, C.J.,AFC[5] |
| Bidi (PI157961, 1947) | PI 157961 TR06ID SD | *Triticum turgidum* L. subsp. *durum* (Desf.) van Slageren | Italy, Sicilia | Bidi | SSG[6] |
| Mahmoudi (PI189779, 1950) | PI 189779 TR04ID SD | *Triticum turgidum* L. subsp. *durum* (Desf.) van Slageren | Tunisia | Mahmoudi | BSV[2] |
| Mahmoudi (PI352413, 1969) | PI 352413 TR03ID SD | *Triticum turgidum* L. subsp. *durum* (Desf.) van Slageren | Tunisia | Mahmoudi | Ingold, M., FARS[6] |
| Mahmoudi (PI41045, 1915) | PI 41045 TR11ID SD | *Triticum turgidum* L. subsp. *durum* (Desf.) van Slageren | Tunisia | Mahmoudi | Guillochon, L., JET[7] |
| Mahmoudi (PI41046, 1915) | PI 41046 TR16ID SD | *Triticum turgidum* L. subsp. *durum* (Desf.) van Slageren | Tunisia | Mahmoudi Ag | Guillochon, L., JET[7] |
| Chili (PI534335, 1976) | PI 534335 TR09ID SD | *Triticum turgidum* L. subsp. *durum* (Desf.) van Slageren | Tunisia, Kebili | Chili | IBPGA[8], FAO in 1976 |
| Chili (PI534336, 1976) | PI 534336 TR09ID SD | *Triticum turgidum* L. subsp. *durum* (Desf.) van Slageren | Tunisia | Chili | IBPGA[8], FAO in 1976 |
| Chili (PI534342, 1976) | PI 534342 TR09ID SD | *Triticum turgidum* L. subsp. *durum* (Desf.) van Slageren | Tunisia | Chili | IBPGA[8], FAO in 1976 |
| Mahmoudi (PI534343, 1976) | PI 534343 TR04ID SD | *Triticum turgidum* L. subsp. *durum* (Desf.) van Slageren | Tunisia | Mahmoudi | CNR[9] in 1976 |

| | | | | | |
|---|---|---|---|---|---|
| Chili (PI534347, 1976) | PI 534347 TR13ID SD | *Triticum turgidum* L. subsp. *durum* (Desf.) van Slageren | Tunisia | Chili | CNR[9] in 1976 |
| Chili (PI534348, 1976) | PI 534348 TR13ID SD | *Triticum turgidum* L. subsp. *durum* (Desf.) van Slageren | Tunisia | Chili | CNR[9] in 1976 |
| Chili (PI534351, 1976) | PI 534351 TR04ID SD | *Triticum turgidum* L. subsp. *durum* (Desf.) van Slageren | Tunisia | Chili | CNR[9] in 1976 |
| Chili (PI534359, 1976) | PI 534359 TR04ID SD | *Triticum turgidum* L. subsp. *durum* (Desf.) van Slageren | Tunisia | Chili | CNR[9] in 1976 |
| Bidi (PI534469, 1976) | PI 534469 TR04ID SD | *Triticum turgidum* L. subsp. *durum* (Desf.) van Slageren | Algeria | Bidi 17 | CNR[9] in 1976 |
| Mahmoudi (PI55538, 1922) | PI 55538 TR03ID SD | *Triticum turgidum* L. subsp. *durum* (Desf.) van Slageren | Tunisia | Mahmoudi AC 3 | Boeuf, F., BSV [2] |
| Mahmoudi (PI55539, 1922) | PI 55539 TR03ID SD | *Triticum turgidum* L. subsp. *durum* (Desf.) van Slageren | Tunisia | Mahmoudi AP 5 | Boeuf, F., BSV[2] |
| Chili (PI576722, 1976) | PI 576722 TR10ID SD | *Triticum turgidum* L. subsp. *durum* (Desf.) van Slageren | Tunisia | Chili | IBPGA[8], FAO in 1976 |
| Bidi (PI576735, 1976) | PI 576735 TR10ID SD | *Triticum turgidum* L. subsp. *durum* (Desf.) van Slageren | Algeria | Bidi 17 | IBPGA[8], FAO in 1976 |
| Bidi (PI576736, 1976) | PI 576736 TR10ID SD | *Triticum turgidum* L. subsp. *durum* (Desf.) van Slageren | Algeria | Bidi 17 | IBPGA[8], FAO in 1976 |
| Bidi (PI576791, 1976) | PI 576791 TR10ID SD | *Triticum turgidum* L. subsp. *durum* (Desf.) van Slageren | Algeria | Bidi 17 | IBPGA[8], FAO in 1976 |
| Mahmoudi (PI7792, 1901) | PI 7792 TR03ID SD | *Triticum turgidum* L. subsp. *durum* (Desf.) van Slageren | Algeria, Setif | Mahmoudi | Fairchild, David Grandison, USDA |

1: UN: University of Nebraska. 2: BSV: Botanical Service of Tunis, Tunisia. 3: ICMEW: Institut Colonial Marseillais Exposition of Wheat. 4: IGA: Inspector General of Agriculture. 5: AFC: American Food Council. 6: SSG: Stazione Sperimentale di Granicoltura. 7: JET: Jardin d'Essais of Tunis, Tunisia. 8: IBPGA: International Board for Plant Generic Ressources, AGPC, FAO, Italy. 9: CNR: Consiglio Nazionale delle Ricerche.

**Table ESM3. Details on the agro-morphological descriptors.**

| Traits | Description | Abbreviation in the study | Source | Descriptor | Codes |
|---|---|---|---|---|---|
| Heading date (Zadoks 50-52) | Regular observations were made to detect the date on which the first spikelet is visible on the spikes of 50% of the plants. Classification was done using the equivalence method. | HS | | Very early | 1 |
| | | | | Early | 3 |
| | | | | Medium | 5 |
| | | | | Late | 7 |
| | | | | Very late | 9 |
| Plant height (Zadoks 75-92) | The height of the plant, expressed in cm, is taken at maturity, from the base of the plant to the extremity of the awns. Classification were done using the equivalence method. | HI | | Very short | 1 |
| | | | | Short | 3 |
| | | | | Medium | 5 |
| | | | | Long | 7 |
| | | | | Very Long | 9 |
| Spike Shape | Note the shape of the spike in profile view. It should be noted that some spike have different shapes than the UPOV scale does not take them into account. | SS | UPOV | Tapering | 1 |
| | | | UPOV | Parallel sided | 2 |
| | | | Personal Observation | Long cylindrical | 2A |
| | | | Personal Observation | Stunted | 2B |
| | | | UPOV | Semi-clavate | 3 |
| | | | UPOV | Clavate | 4 |
| | | | UPOV | Fusiform | 5 |
| Spike Colour | Attribute a colour gradient of each spike after observing the maximum of spikes. | SC | UPOV | White | 1 |
| | | | | Slightly coloured | 2 |
| | | | | Strongly coloured | 3 |
| Spike Density | A count of the number of spikelets in 10 cm of spikes was carried out. | SD | IPGRI | Very lax < 16 | 1 |
| | | | | Lax 16- 25 | 2 |
| | | | | Medium 25.1 -30 | 3 |
| | | | | Dense 30.1-40 | 4 |
| | | | | Very dense >40 | 5 |
| Length of Awns in relation to Spike | Measurement of awn and spike lengths | LAS | UPOV | Shorter | 1 |
| | | | | Equal | 2 |
| | | | | Longer | 3 |
| | | HAR | UPOV | Absent or very weak | 1 |

| | | | | | |
|---|---|---|---|---|---|
| Hairiness of margin of first rachis segment | An observation of the margin of first rachis segment was made to see if the hair is absent or present and to estimate its abundance. | | | Weak | 2 |
| | | | | Medium | 3 |
| | | | | Strong | 4 |
| | | | | Very strong | 5 |
| Length of Spike Without Awns | Measurement of spikes length without counting awns. Classification was done using the equivalence method. | LSWA | UPOV | Very short | 1 |
| | | | | Short | 3 |
| | | | | Medium | 5 |
| | | | | Long | 7 |
| | | | | Very long | 9 |
| Length of Awns | Measurement of awns length. Classification was done using the equivalence method. | LA | | Very short | 1 |
| | | | | Short | 3 |
| | | | | Medium | 5 |
| | | | | Long | 7 |
| | | | | Very long | 9 |
| Awn Colour | Attribute a colour of each awn after observing the maximum of awn spikes. | AC | UPOV | Whitish | 1 |
| | | | | Light brown | 2 |
| | | | | Brown | 3 |
| | | | | Black | 4 |
| Anthocyanin colouration of Awns | Attribute a gradient of anthocyanin colouration of each awn after observing the maximum of awn spikes. | PgA | UPOV | Absent or very weak | 1 |
| | | | | Weak | 3 |
| | | | | Medium | 5 |
| | | | | Strong | 7 |
| | | | | Very strong | 9 |
| Distribution of Awns | An observation of the distribution of awns along the spike | DtA | UPOV | Awnless | 1 |
| | | | | Tip only | 2 |
| | | | | Upper half | 3 |
| | | | | Whole length | 4 |
| Number of Spikelet by Spike | A count of the number of spikelets per spike. Classification was done using the equivalence method. | NSS | IPGRI | Very weak | 1 |
| | | | | Weak | 2 |
| | | | | Medium | 3 |
| | | | | Strong | 4 |
| | | | | Very strong | 5 |
| Colour of Grain | Attribute a colour of each lot of grain after observing the maximum of spikes using a colour chart. | CG | | Ral 1011 | 1 |
| | | | | Ral 8024 | 2 |
| | | | | Ral 8003 | 3 |

| | | | | | |
|---|---|---|---|---|---|
| | | | | Ral 8001 | 4 |
| Shape of Grain | Attribute a form of each lot of grain after observing the maximum of grain lots. | SG | UPOV | Ovoid | 3 |
| | | | | Semi-elongated | 5 |
| | | | | Elongated | 7 |
| Thousand Grain Weight | A count of the number of grains by spike and a weighing of the grains by spike. Calculation of weight of 1 grain ant then of the Thousand grain. Classification was done using the equivalence method. | TGW | | Very weak | 1 |
| | | | | Weak | 2 |
| | | | | Medium | 3 |
| | | | | Strong | 4 |
| | | | | Very strong | 5 |
| Area Under Disease Progression Curve | Scoring of *Septoria tritici* blotch at two different dates using Saari and Prescott's "double digit" method (Saari & Prescott, 1975).<br>Area Under the disease progression curve (AUDPC) was calculated from the percentage of disease severity (DS) at both observation dates (Sharma and Duveiller, 2007; Das & al., 1992). Classification was done using the arithmetic progression method. | AUDPC | | Very resistant | 1 |
| | | | | Resistant | 2 |
| | | | | Moderately resistant | 3 |
| | | | | Susceptible | 4 |
| | | | | Very susceptible | 5 |
